# Genome-Wide Annotation of Promoter-Enhancer Interactions via Chromatin Loops in the Hybrid Rat Diversity Panel

**DOI:** 10.1101/2025.09.10.675455

**Authors:** Panjun Kim, Burt M. Sharp, Robert W. Williams, Hao Chen

## Abstract

Despite the prominence of laboratory rats in behavioral neuroscience and complex trait genetics, a significant gap exists in the functional rat genomics database that explains the regulation of gene expression at the genome level. To address this, we analyzed genome-wide Hi-C data from the frontal cortex of ten strains in the Hybrid Rat Diversity Panel. While originally generated to improve the rat genome assembly, these data provided a unique opportunity to characterize the regulatory landscape of the adult rat brain. We identified an average of 5,899 ± 1,997 (STD) loops per sample and integrated these with over 3 million curated CTCF binding sites, which serve as architectural anchors for chromatin loops. Our multi-stage filtering workflow identified 15,085 unique, high-confidence regulatory interactions throughout the genome. As a validation of our approach, we observed that genes with the highest loop counts, including *Foxo1, Gja1*, and *Spry2*, were significantly enriched for developmental processes, reflecting the role of 3D genome structure in maintaining adult neuronal identity. The resulting resource addresses a critical deficiency in rat functional genomics and provides a foundation for dissecting the genetic architecture of complex traits.

## INTRODUCTION

The spatiotemporal control of gene transcription is fundamental to the development and maintenance of complex biological systems (1). This regulation is primarily mediated by distal cis-regulatory elements, such as enhancers, which integrate physiological and environmental signals to drive cell-type-specific transcriptional profiles (1, 2). In the central nervous system, these regulatory mechanisms underlie the expression of complex behavioral phenotypes (3–5). Genetic variants in these non-coding regions are frequently associated with phenotypic diversity and susceptibility to psychiatric conditions (6–8). Identifying the genes regulated by these elements is essential for characterizing the molecular mechanisms that link genotype to phenotype.

The laboratory rat (*Rattus norvegicus*) is a prominent model for biomedical research, offering a physiological complexity and behavioral repertoire, including reward processing and social behavior, that parallels human conditions more closely than other rodent models (9–11). Although the rat reference genome has recently reached a quality comparable to human and mouse standards (12–14), its functional annotation remains significantly underdeveloped (15). This gap is underscored by the exhaustive functional mapping available for other major mammalian models through initiatives such as ENCODE and Roadmap Epigenomics (15). No such comprehensive resource exists for the rat, leaving much of its regulatory landscape undefined. Furthermore, because regulatory relationships are highly tissue- and cell-type specific, the lack of data from relevant brain regions hinders the interpretation of how genetic variation influences physiological and behavioral outcomes (8).

Mapping the physical interactions between distal enhancers and their target promoters is a critical step toward overcoming this limitation. The three-dimensional (3D) organization of the genome facilitates these contacts, often through chromatin loops organized within topologically associating domains (TADs) (2, 16, 17). These architectural features are typically anchored by proteins such as the CCCTC-binding factor (CTCF) and cohesin, which bring distant enhancers, where transcription factors bind, into proximity with gene promoters to regulate transcription (1, 18). High-throughput chromosome conformation capture (Hi-C) provides a robust methodology for resolving these interactions at a genome-wide scale, enabling the identification of promoter-enhancer (P-E) loops that control gene expression (1, 18). However, while high-resolution mapping of enhancer-promoter contacts has been extensively characterized in various human and mouse cell types (1), equivalent comprehensive 3D genomic and epigenomic annotations for the rat lag significantly behind those of other major mammalian models (11, 15).

In this study, we use Hi-C to delineate the 3D regulatory architecture of the rat frontal cortex, a region central to executive function and behavioral regulation. We analyzed data from ten inbred strains within the Hybrid Rat Diversity Panel (HRDP), a reference population designed for high genetic and phenotypic diversity (19, 20). These data, originally generated for *de novo* genome assembly of individual HRDP strains, represent a high-resolution resource that we repurposed to characterize the regulatory landscape of the rat brain. By integrating these contact maps with over 3 million CTCF binding sites predicted using FIMO, we identified 15,085 high-confidence regulatory interactions across the rat genome, with each interaction assigned to a specific target gene.

The validity of these gene-specific loops is strongly supported by the finding that genes with the highest loop density, including *Foxo1, Gja1*, and *Spry2*, are significantly enriched for developmental processes essential for maintaining adult neuronal identity. This enrichment is consistent with the role of 3D genome architecture in preserving transcriptional states required for mature neuronal function. This resource addresses a critical deficiency in rat functional genomics, providing a foundational tool for the research community to investigate the molecular drivers of behavioral and physiological diversity.

## RESULTS

### Characterizing loops from 10 Hi-C rat samples

The production of these sequencing data has been described in detail (21), and all data have been deposited to the NIH Short Read Archive under accession number PRJNA1197090. Briefly, Arima Hi-C libraries were extracted from the frontal cortex of 10 rats, each a unique strain or F1 hybrid: SHR/OlaIpcv, BN-Lx/Cub, BXH6/Cub, HXB2/Ipcv, HXB10/Ipcv, HXB23/Ipcv, HXB31/Ipcv, LE/Stm, F344/Stm, and SHR/OlaIpcv x BN/NHsdMcwi F1, selected to maximize genetic distances based on a phylogenetic tree we generated before (14). Data were processed using the Juicer pipeline (v1.6) (22), aligned to the mRatBN7.2/rn7 reference genome. On average, 570.2 million paired-end reads were generated per sample, although LE/Stm and F344/Stm samples yielded fewer reads (21). Figure 1a summarizes the distribution of several categories of reads as proportions of the total read pairs. Unalignable reads, which include chimeric, ambiguous, and unmapped categories, accounted for an average of 11.4% of the total sequenced read pairs. Among these, unmapped reads represented only 1.3%. The overall proportion of unalignable reads was the lowest among all read categories and showed relatively low variability across samples (STD = 2.1%). PCR and optical duplicates accounted for 21.4% of the reads. Notably, samples such as LE/Stm (8.0%) and F344/Stm (8.6%) showed markedly lower duplication rates. The variation in duplication rates across samples (STD = 11.0%) most likely reflects technical differences. Unique reads constituted the largest proportion of total sequenced read pairs across all samples (67 ± 10.0%), but the extent varied considerably among samples. We retained uniquely mapped reads with a mapping quality threshold of 30 (i.e., < 0.1% chance of misalignment) to ensure reliable Hi-C contact inference.

**Figure 1.**
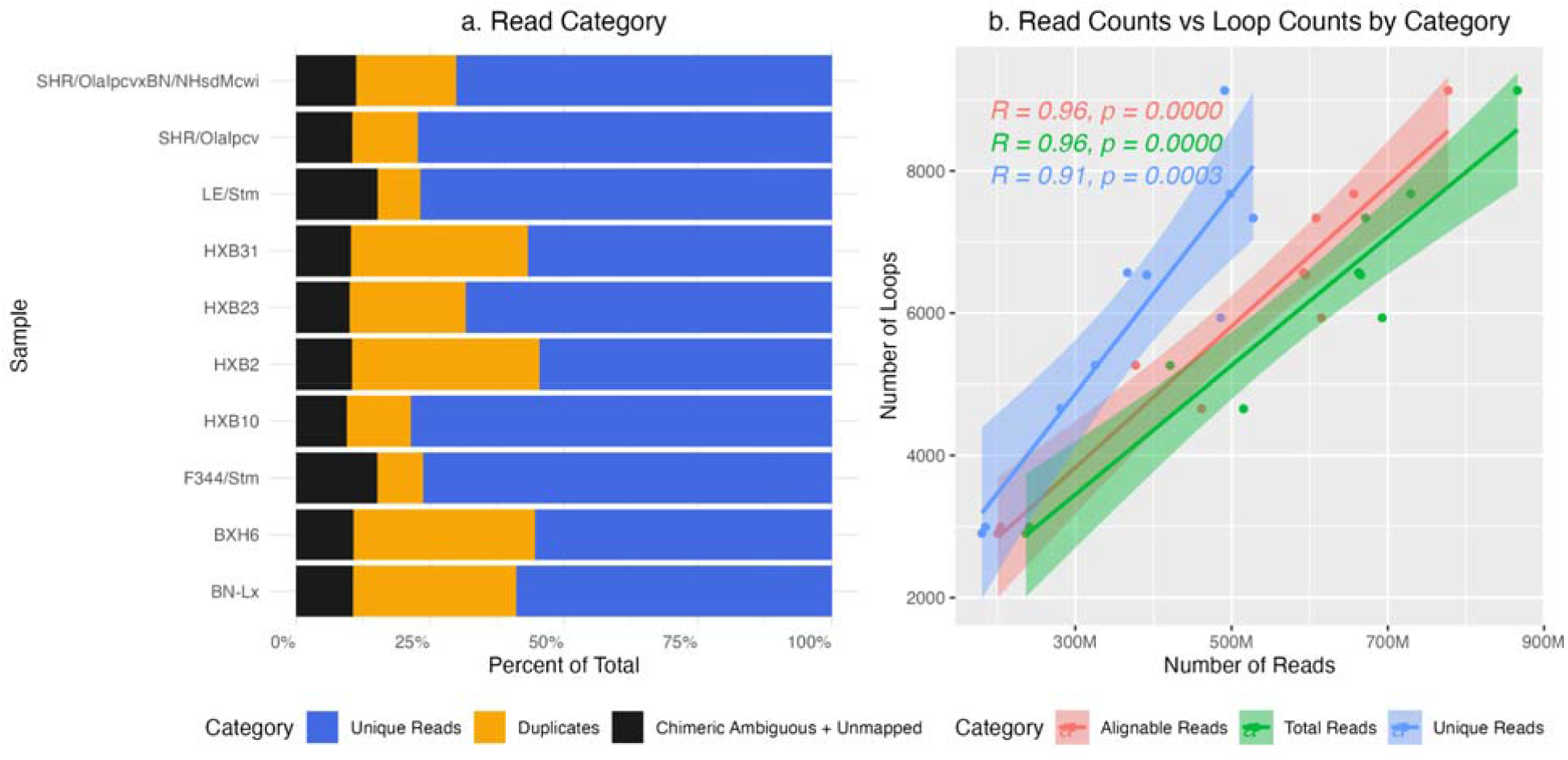
Depth of unique reads determines the number of detected loops. **a**. Proportions of unique, duplicate, and chimeric/unmapped reads in Hi-C data across 10 samples. Hi-C sequencing reads from 10 samples were processed using the Juicer pipeline (v1.6) and categorized into unique reads, PCR/optical duplicates, and unalignable reads (including chimeric ambiguous, and unmapped). Unique reads comprised the majority across most samples. **b**. Positive correlation between loop counts and sequencing depth across 10 samples.

We further processed the Hi-C sequencing data using HiCCUPS (18) from the Juicer pipeline to annotate loop structures. In total, 58,992 loops were identified across the 10 samples, with per-sample counts ranging from 2,903 to 9,131 (Table S1). To evaluate the relationship between sequencing metrics and chromatin loop detection, we compared the number of loops identified per sample as a function of three sequencing depth metrics: total reads, unique reads, and alignable reads (Figure 1b). Loop counts were strongly correlated with sequencing depth for all three metrics examined (total reads, alignable reads, and uniquely mapped reads), with Pearson correlation coefficient ranging from 0.91 to 0.96 (all p < 0.001; Figure 1b), indicating that sequencing depth is a primary determinant in loop detection. In addition, several other QC metrics (Figure S1), such as the number of Hi-C contacts and the proportion of non-chimeric alignable reads, also showed strong positive associations with loop count. These results underscore the primary role of sequencing depth and library complexity in identifying chromatin structures.

We investigated how loop counts varied across three loop calling resolutions (5K, 10K, and 25K), considering all chromosomes (Figure 2). Overall, the number of annotated loops generally increased with broader resolution, particularly from 5K to 10K, with more modest gains observed between 10K and 25K. An exception to this trend was the Y chromosome, which showed a decrease in loop counts at 25K. Across the genome, the number of detected loops was strongly and positively correlated with chromosome length (r = 0.88, p = 5.47 × 10□□) as expected, reflecting more chromatin regulatory interactions in larger chromosomes.

**Figure 2.**
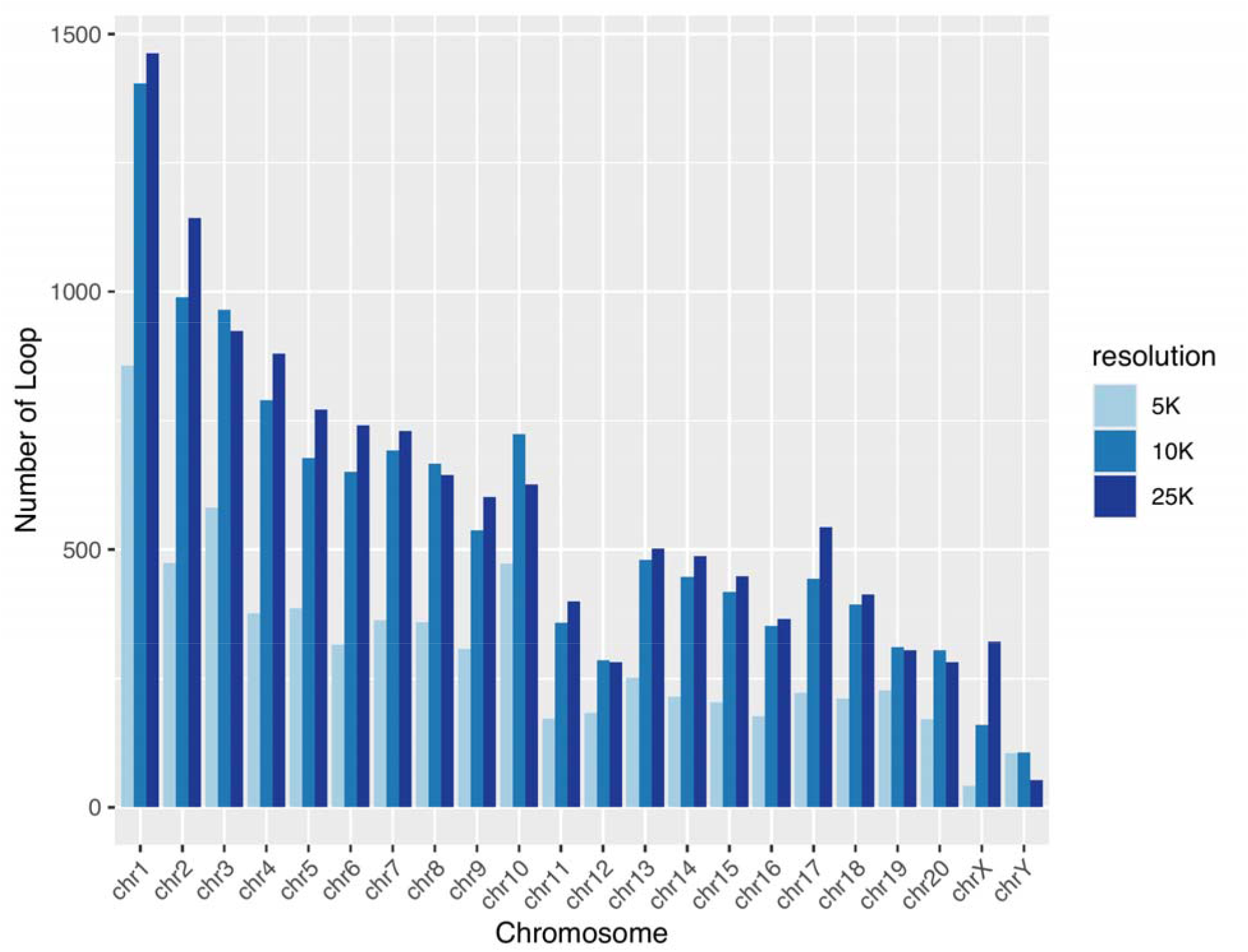
Genome-wide distribution of loop counts detected at three resolutions. Loops were annotated on each chromosome at 5K, 10K, and 25K resolutions using HiCCUPS. In general, higher resolution (5K) yielded fewer detected loops. Larger chromosomes tend to have more loops, reflecting a greater number of regulatory interactions.

We then examined loops that were detected in multiple samples at each resolution. Figure 3a summarizes the mean number and variability of loops shared (i.e., have identical anchoring positions) between any two samples at each resolution. Lower resolution (25K) yielded a greater number of shared loops, accompanied by reduced variability. Specifically, the average number of shared loops increased from 774 ± 468 at 5K resolution to 1,379 ± 420 at 10K and 1,853 ± 225 at 25K. In addition, there was substantial variability in the total number of loops per sample (Figure 3b), with HXB31 showing the highest count (9131), and 54.6% classified as shared loops compared to 45.4% unique loops. Conversely, F344/Stm, which had the lowest total loop count (2903), demonstrates 74.8% shared loops and 25.2% unique. Notably, samples with fewer detected loops exhibited a higher proportion of shared loops, likely due to relatively low sequencing depth in these samples.

**Figure 3.**
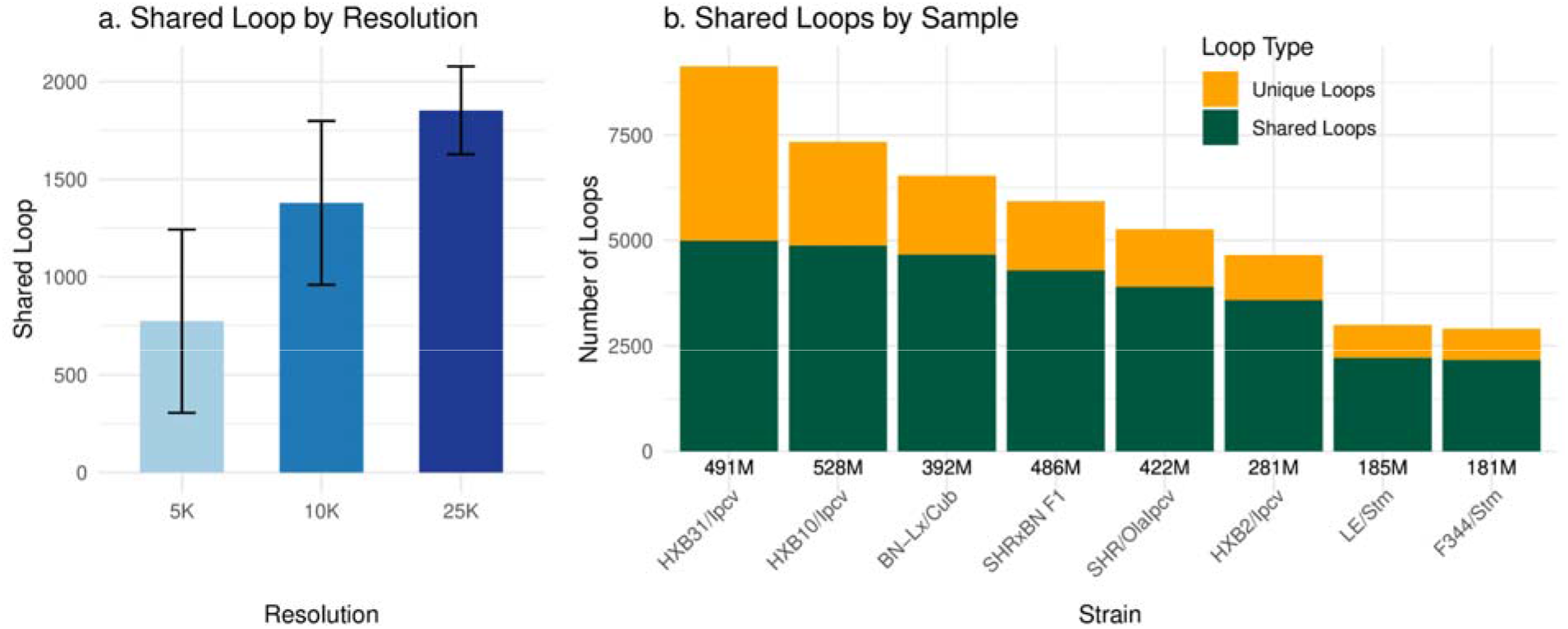
The majority of the loops are shared between samples. **a.** Number of shared chromatin loops between any pairs of samples at 5K, 10K, and 25K resolutions. The number of shared loops was greater, and variability was lower at lower (i.e., 25K) detection resolution. Error bars represent standard deviations. **b**. Distribution of unique and shared loops across rat samples. Chromatin loops are classified as either unique (sample-specific) or shared (observed in multiple samples) across each of the ten samples. Samples with greater sequencing depth (e.g. HXB31 and HXB23) have more loops detected and higher proportion of unique loops, compared to samples with lower number of sequencing depth (e.g., LE/Stm and F344/Stm).

After combining loops from all three resolutions (Figure S2), we observed a high degree of sharing, suggesting a conserved chromatin topology among these related strains. In contrast, fewer connections in other strains reflect greater biological divergence, combined with artifacts of technical variability. Figure S3 further quantifies pairwise loop sharing across three resolutions (5K, 10K, and 25K) using heatmaps. At 5K resolution, F344/Stm, HXB2, and LE/Stm exhibited a relatively higher proportion of shared loops compared to higher resolutions. This pattern may, in part, result from lower total loop counts in these samples. Together, these data reveal both conserved and strain-specific features of chromatin looping among a diverse collection of rat genomes.

### Genome-wide annotation of CTCF binding sites

To further validate the functional relevance of these loops, we examined the genomic features that are critical for P-E interactions. CTCF plays a critical role in the formation of chromatin loops by acting as a boundary element that binds to specific DNA motifs. Accordingly, mapping the positional distribution of CTCF motifs provides a framework for identifying loops that may contribute to transcriptional regulation. Using 100 CTCF motifs curated from six databases (Table S2), we identified 6,551,641 potential CTCF binding sites across the rat genome using FIMO (23). The ideogram in Figure 4a illustrates the genomic distribution of CTCF elements across the rat genome. CTCF binding sites were not uniformly distributed across the chromosomes. We found chromosome length was not correlated with CTCF density per Mb (Pearson r = -0.202, p = 0.367). In contrast, CTCF density was strongly correlated with gene density (Figure 4b-4d). Using NCBI annotations, CTCF density was significantly correlated with the density of all genes (Pearson r = 0.922, p = 1.03e-09), protein-coding genes (r = 0.877, p = 8.51e-08), and lncRNA genes (r = 0.891, p = 2.66e-08). Similar correlations were observed using Ensembl annotations for all genes (r = 0.910, p = 4.56e-09), protein-coding genes (r = 0.872, p = 1.24e-07), and lncRNA genes (r = 0.796, p = 9.32e-06).

**Figure 4.**
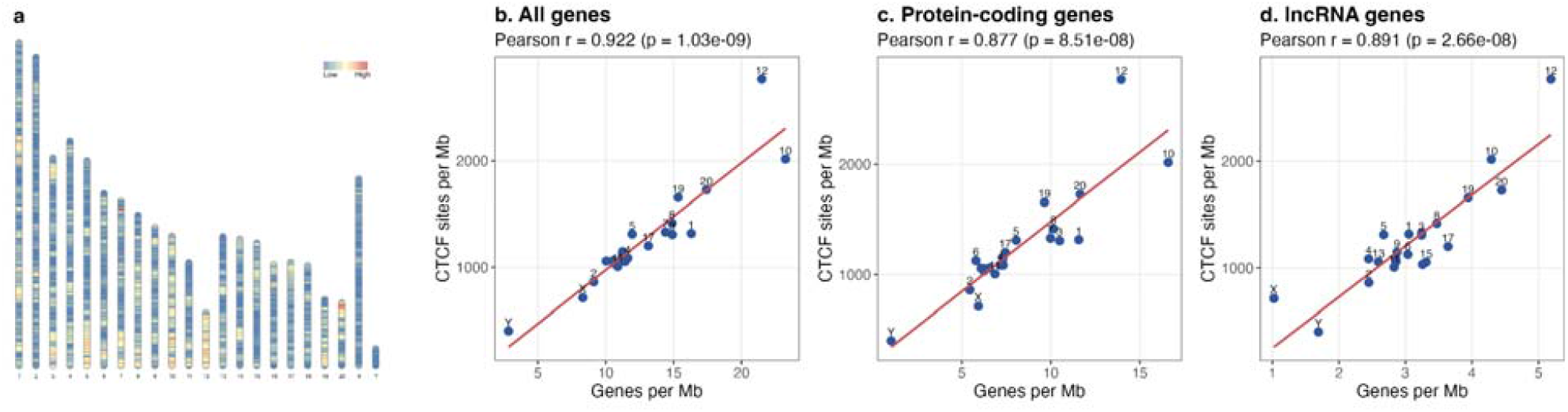
Chromosomal distribution of CTCF binding site density and its correlation with gene density across the rat genome. **a.** Genome-wide distribution of CTCF binding sites across all rat chromosomes (mRatBN7.2 assembly). Each vertical bar represents a chromosome, with color-coded segments reflecting the density of CTCF binding sites within non-overlapping 1 Mb genomic bins. The heatmap scale ranges from low (blue, minimum number of 0 CTCF in a bin) to high (red, maximum number of 5711 CTCF in a bin) density, allowing for visual comparison of CTCF abundance across different chromosomal regions. **b–d.** Chromosome-level correlation between gene density (genes per Mb) and CTCF binding site density (CTCF sites per Mb) using NCBI gene annotations. Each data point (blue dots) represents a single chromosome. **b.** All genes (n = 42,049), **c.** protein-coding genes (n = 21,944), and **d.** lncRNA genes (n = 7,815). Pearson correlation coefficients and p-values are shown in each panel. Linear regression fits (red lines) highlight the positive association between gene density and CTCF density.

### Positional patterns of CTCF, TSS, and promoter near loop anchors

In addition to CTCF binding sites, we also used 42,925 Ensembl start-codon annotations and 12,601 experimentally validated promoter coordinates. After filtering and removing redundant entries based on identical coordinates (chromosome, start, and end positions), we retained 3,191,859 unique CTCF binding sites, 21,725 distinct TSS, and 12,529 unique promoter regions for downstream analyses. We next examined the distribution of CTCF motif binding sites, TSS, and promoter regions in the vicinity of chromatin loops. We used a final set of 31,019 loops shorter than 2Mb (18, 24). For the positional distribution analysis, we analyzed a slightly reduced set of 30,928 loops, excluding the 91 loops located at the end of a chromosome that exceeded the 3L flanking region definition.

Figure 5 and Figure S4 present density plots and histograms, respectively, illustrating the relative positions of CTCF (**a**), TSS (**b**), and promoters (**c**) across all loops. Each plot spans flanking regions equivalent to one loop length on either side of the anchor. The density plots in Figure 5 are stratified by three different resolutions (5K, 10K, and 25K). The bimodal pattern is evident across all resolutions, with distinct peaks showing enrichment of CTCF (Figure 5a) occurring symmetrically at positions flanking the loop region (approximately at 0 and +1, which represent the start and end of the loop, after normalization of loop length). The pronounced enrichment at loop boundaries underscores the architectural role of CTCF in anchoring chromatin loops, a feature that is consistently maintained across resolutions. While this pattern remains stable, peak intensity tends to increase with higher resolution. A bimodal enrichment of TSS (b) and promoter (c) is also evident at loop boundaries; however, the peaks in these figures appear broader, and resolution-dependent differences in peak height are less distinct.

**Figure 5.**
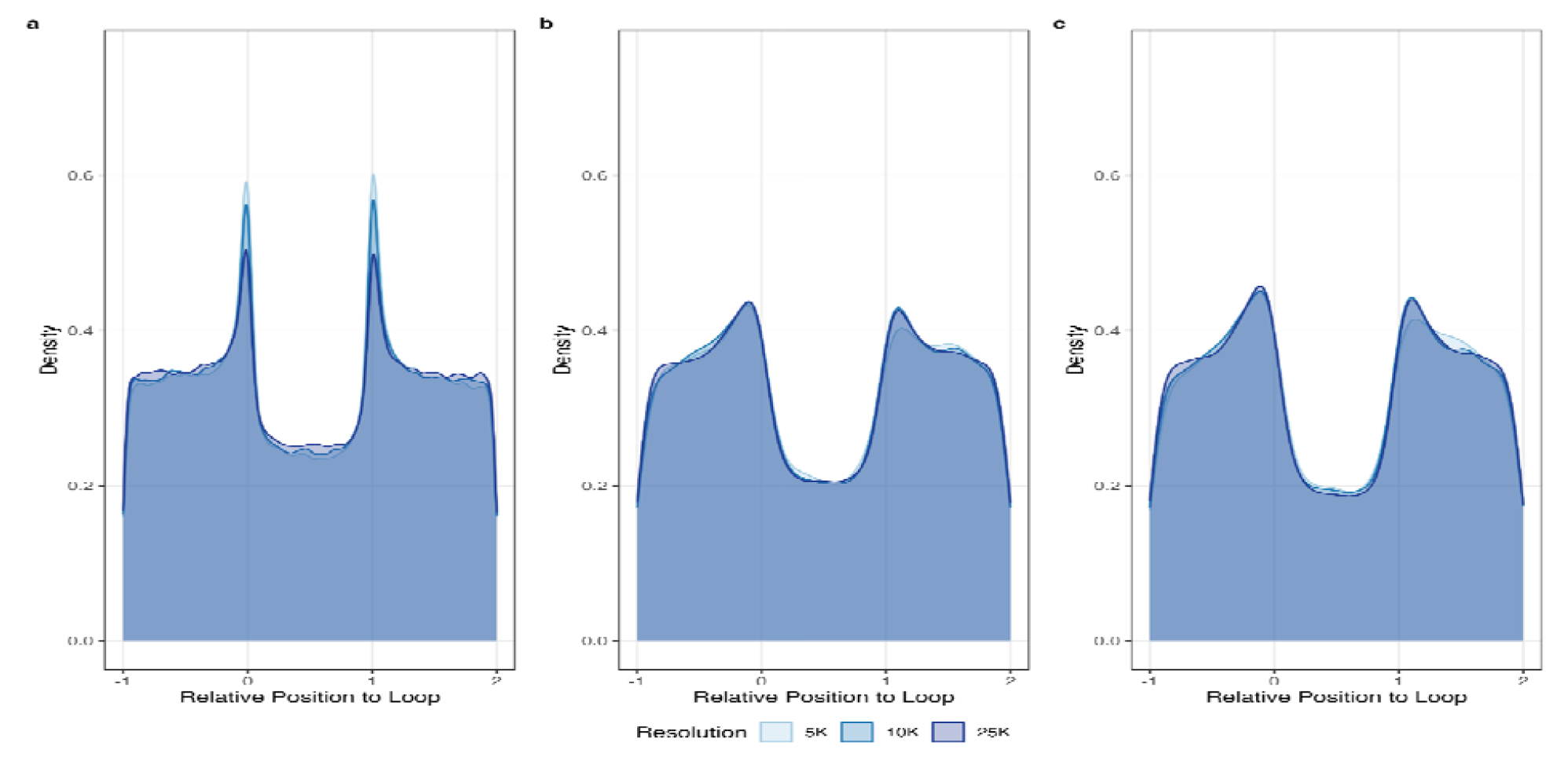
Density plots for positional distribution of CTCF binding sites, TSS, and promoters relative to chromatin loops. Each plot spans ±1 loop length from the loop boundaries, with resolution-specific curves overlaid. **a**. CTCF binding sites exhibit distinct peaks at both loop anchors (x = 0 and 1) across all resolutions, indicating consistent enrichment at loop boundaries regardless of resolution. **b**. TSS density displays an enrichment near loop anchors with lower and broader amplitude compared to CTCF. **c**. The distribution of the promoter also displays a similar profile near both anchors resembling the pattern observed for TSS.

At the level of each chromosome, density plots and histograms for the relative location of CTCF, TSS, and promoter elements showed more diverse distributional patterns around chromatin loop anchors. These patterns ranged from distinct, symmetric peaks at both loop anchor locations to asymmetric or weakly defined profiles, indicating variability in their spatial positioning and enrichment (Figure S5, S6, and S7).

Informed by the positional distribution of CTCF motifs, TSS, and promoter elements, we examined genomic features near the anchors of chromatin loops to establish a threshold for selecting loops that mediate P-E interaction. We used the final set of 31,019 loops shorter than 2 Mb for this and all subsequent structural filtering analyses. For CTCF, we quantified the number of CTCF motif binding sites present in both anchors of the loops defined by HiCCUPS. The distribution of CTCF binding site counts in the anchors showed a distinct bimodal pattern across all detection resolutions (**Figure 6**). A prominent peak corresponded to the majority of loops, centered around approximately 32 CTCF motifs per anchor for loops detected at 5K or 10K resolution, and approximately 42 motifs for those detected at 25K resolution. In addition, a minor peak with 1 or 2 CTCF sites likely arose from artifacts of the loop detection method. Regardless of resolution, the trough between these two populations occurred at approximately 6 CTCF sites and served as a data-driven threshold to distinguish likely functional loops from those likely falsely detected by HiCCUPS.

**Figure 6.**
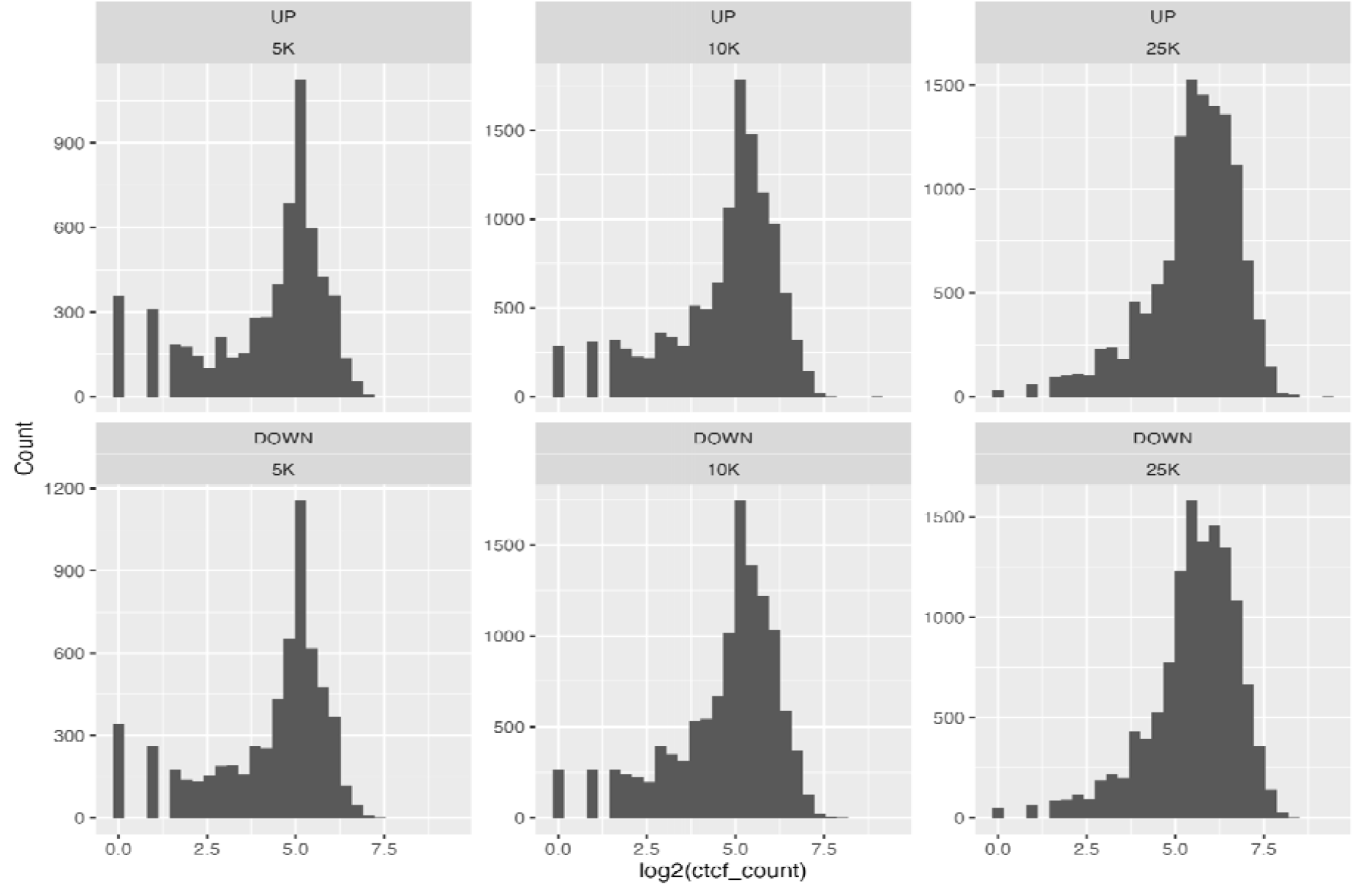
Distribution of CTCF binding site counts at loop anchors by direction and resolution. Histograms showing the distribution of CTCF binding site counts (log□-transformed) at loop anchors, stratified by loop resolution (5K, 10K, and 25K) and direction (upstream vs. downstream), where upstream and downstream are labeled as “UP” and “DOWN” respectively in the plot. Each panel represents a specific combination of direction and resolution, illustrating how CTCF occupancy varies across the genomic scale and loop orientation. A distinct bimodal distribution is evident across all panels. The first peak, consisting of loops with a low number of CTCF domains (log□ (ctcf_count) < 2.5), is likely noise resulting from the loop detection method. Of the 31,019 loops analyzed, only 111 lacked detectable CTCF binding sites at both the upstream and downstream anchors.

### Identifying loops mediating P-E interaction

We conducted extensive annotation and filtering steps, as shown in Figure 7, to identify chromatin loops that are likely to mediate P-E interactions. These filtering steps are based on the assumption that one loop anchor is located near the start of a gene, that is, the promoter or TSS, while the other anchor corresponds to a regulatory element, such as an enhancer, and that the loop is flanked by strong CTCF signals.

**Figure 7.**
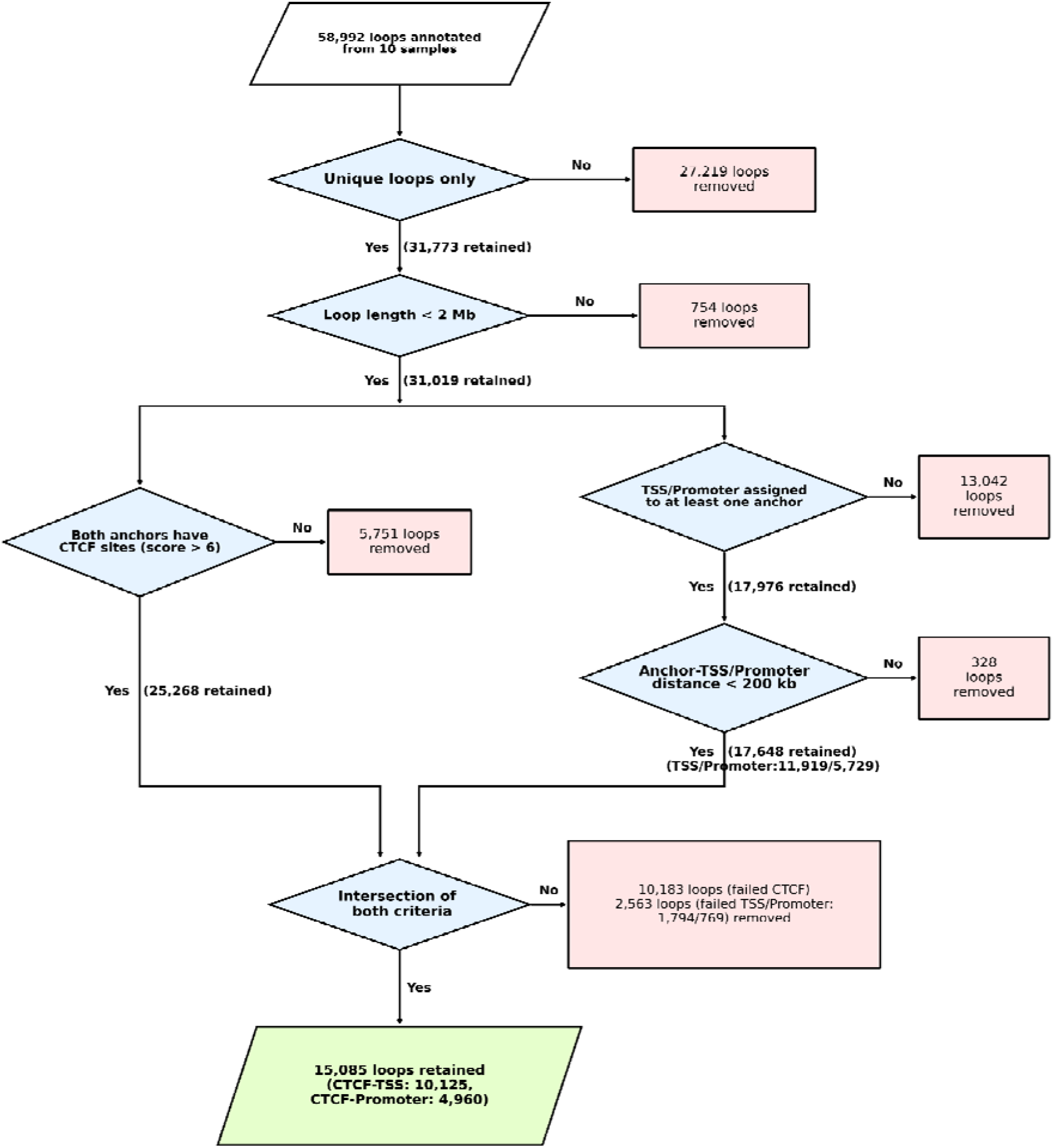
Workflow for the identification of high-confidence gene regulatory chromatin loops. The filtering pipeline began with an initial aggregate of 58,992 loops identified across ten biological samples. Duplicate loops were merged to eliminate redundancy (27,219 loops removed), followed by the exclusion of long-range interactions exceeding 2 Mb to prioritize intra-domain contacts (754 loops removed). The remaining loops were subjected to a dual-criteria assessment: (1) Architectural filtering, which required both loop anchors to contain a minimum of six unique CTCF binding sites; and (2) Functional annotation, which selected loops containing a valid TSS or promoter region within 200kb of the anchor-to-feature distance. Only interactions satisfying both the architectural and functional criteria were retained, yielding a final dataset of 15,085 high-confidence loops.

We first assigned the nearest promoter or TSS to each anchor. Only TSS or promoters located on the same side of the loop, defined by the loop midpoint, were considered. After excluding distance-tied cases, we obtained 61,969 anchors with a single TSS or promoter assignment.

Three additional anchors retained two valid assignments each, resulting in a total of 61,975 anchor–regulatory element assignments. Next, we examined the distance between the assigned promoter or TSS and the corresponding anchor. As shown in Figure S8, more than half of the promoters or TSS overlap with the anchors. However, the distance distribution exhibits a long tail. We reasoned that a minority of promoters or TSS located more than 200 kb away from the anchor are unlikely to be biologically relevant. Applying this distance threshold retained 11,919 loops associated with TSS and 5,729 loops associated with promoter regions, resulting in a total of 17,648 loops selected for downstream analysis.

We next filtered these loops for CTCF signals. Applying the threshold of at least six CTCF motifs in both anchors identified 25,268 loops, which represents 81.5% of the total 31,019 loops analyzed. The intersection of these two sets was then assessed to determine the extent to which CTCF□ anchored loops were also supported by TSS□ or promoter□based assignments. This analysis identified 10,125 loops that satisfied both the TSS and CTCF□ based criteria, and 4,960 loops that satisfied both the promoter and CTCF□based criteria, resulting in a total of 15,085 loops supported by both structural (CTCF) and functional (TSS or promoter) evidence.

The genomic location of the chromatin loops likely mediating P-E interactions, as illustrated in Figure 8a for the entire genome and Figure 8b for Chr 1, showed that they are widely distributed across chromosomes, though some genomic regions are more densely populated than others, while others appear depleted. Circos plots of the loops for other chromosomes are provided in Figure S9.

**Figure 8.**
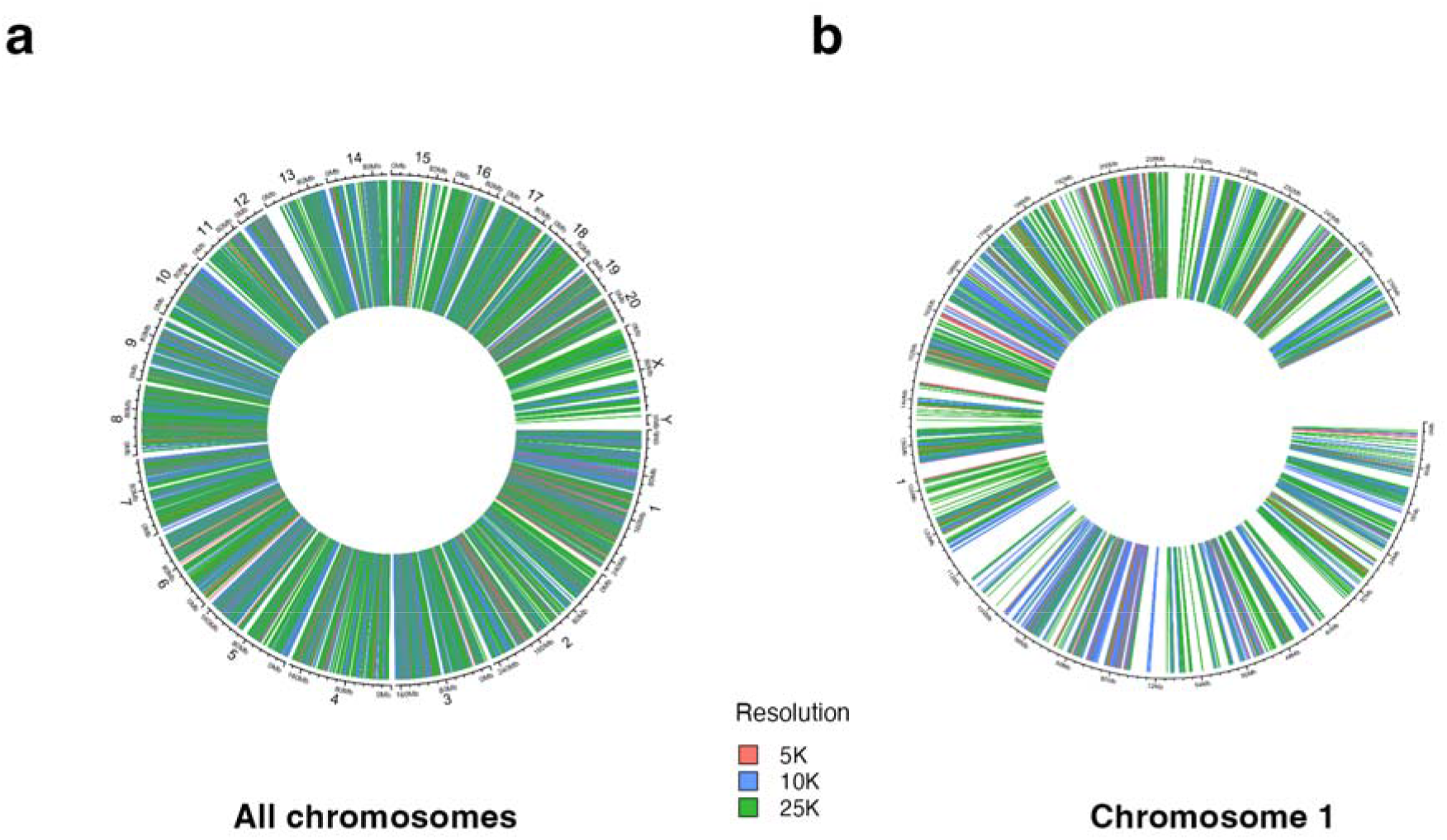
Circos plots of high-confidence chromatin loops across the genome (a) and chromosome 1 (b), stratified by resolution. **a.** genome-wide chromatin loops stratified by resolution (5K, 10K, and 25K). **b.** loops located on chromosome 1. Each arc represents a chromatin loop between two genomic loci, with arc heights scaled logarithmically to reflect their genomic span. These plots illustrate the broad yet non-uniform distribution of loops and enable visual comparison across resolution levels.

We next counted the number of loops each gene is associated with. We found a total of 54 genes associated with 11 or more loops (Table S3). The top three most connected genes, *Foxo1, Gja1*, and *Spry2*, were annotated with 19 loops each. To investigate potential functional convergence among the interaction-rich genes, we performed gene ontology enrichment analysis using g:Profiler (25) on the top 54 genes, corresponding to those with 11 or more interactions (Figure 9). Analysis using the top 25 genes, each with at least 13 interactions, showed similar results. We found a coherent enrichment in embryonic development, morphogenesis, cellular proliferation and other broader developmental growth and stimulus□response functions. Many of these ontological categories are related and contain overlapping genes, such as *Spry2, Foxc1*, and *Tbx3*.

**Figure 9.**
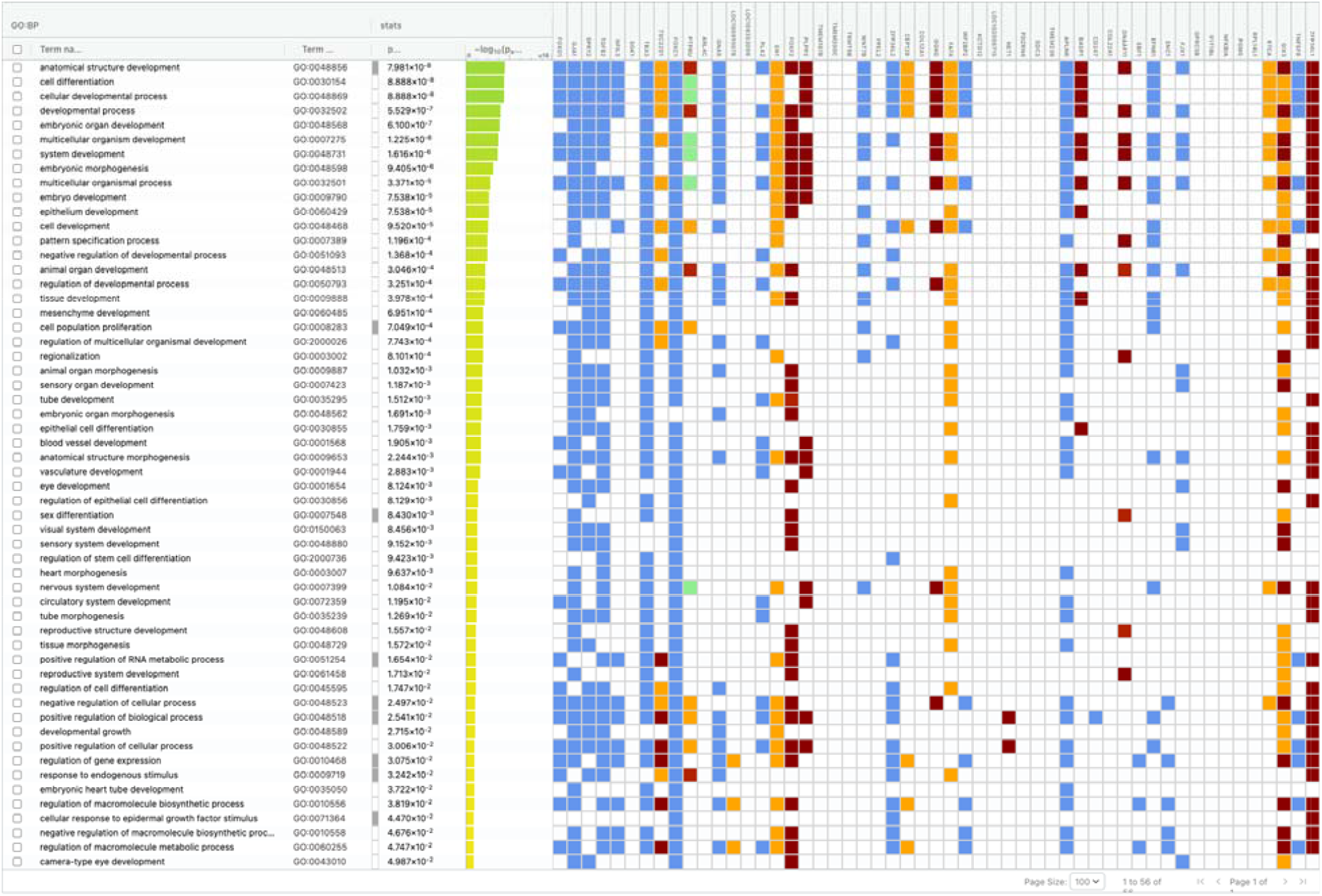
Functional enrichment of top interaction-rich genes reveals developmental and proliferative biological processes. Functional enrichment analysis of the top 54 genes with the highest numbers of loop interactions was performed using g:Profiler. The figure highlights the presence of key developmental and proliferative processes, including embryonic organ development, tissue morphogenesis, and cell population proliferation, across representative gene loci.

## DISCUSSION

We generated a map of chromatin loops for the frontal cortex of ten rat strains from the HRDP. Integrating Hi-C data with genomic annotations identified 15,085 loops with strong likelihood of modulating gene expression. By calling loops across ten diverse genetic backgrounds and merging them into a consensus map, we established a robust multi-strain regulatory resource. This dataset accounts for the inherent genetic diversity of the HRDP, providing a foundation for investigating the 3D landscape of rat genomes and facilitating the dissection of genetic architecture underlying complex behavioral traits.

CTCF is an 11-zinc-finger DNA-binding protein involved in regulating the 3D structure and function of the eukaryotic genome (26, 27). Current models posit that the 3D genome is organized via loop extrusion, a process where cohesin extrudes chromatin until halted by inward-facing CTCF anchors (28, 29). In the absence of experimental CTCF data for the rat genome, we used the FIMO motif scanning tool to predict CTCF binding sites on the mRatBN7.2 reference genome. The predictive accuracy of this method is supported by comparative studies in human and mouse genomes, where FIMO-predicted CTCF sites showed an approximately 80% true positive rate against experimental data (30). To increase the recall of CTCF sites, we included 100 curated CTCF motifs from multiple databases (Table S2). While this approach may increase the number of false positive sites, our analysis of the location of CTCF sites near loop anchors revealed a distinct bimodal distribution as expected (Figure 5). We observed this pattern across all three resolutions, consistent with an architectural role for CTCF in loop formation (18, 31, 32). We also observed that the predicted CTCF binding sites are not uniformly distributed across chromosomes (Figure 4a). Instead, they are associated with gene density, a pattern observed across independent gene annotations and multiple gene classes, such as protein-coding gene and long non-coding gene (Figure 4b-d). This also provides additional evidence that the predicted CTCF binding sites are reliable despite the use of cross-species motifs.

To systematically identify functional loops facilitating P-E interactions, we implemented a multi-stage filtering workflow (Figure 7). We first enriched for structurally stable loops by requiring at least six CTCF binding sites at both anchors based on our data. Under this criterion, 81.5% of the loops were retained. It is possible that this high stringency might have excluded those dynamic, tissue-specific P-E interactions mediated by factors such as the Mediator complex in the absence of dense CTCF clusters (33). We next require that at least one anchor located in the vicinity (i.e. <200 kb) of a promoter or TSS. This distance threshold aligns with established mammalian models where the majority of functional interactions occur within a 200 kb radius of the anchors (34). This step resulted in the exclusion of 13,043 loops, which likely were structural in nature rather than regulatory. In cases where both anchors contained TSS/promoter annotations, we resolved the ambiguity by selecting the element closest to its respective anchor midpoint as the primary target.

The biological relevance of the loops in our dataset is supported by the presence of many genes with a large number of loops (Table S3), such as *Foxo1, Gja1, Spry2, Tgfb2* and *Tbx3*. Most of these genes are involved in embryonic development as shown by GO enrichment analysis (Figure 9). Consistent with the findings that key developmental genes frequently possess ten or more redundant enhancers (35), we infer that the dense loop architectures around these loci correspond to complex multienhancer regulatory landscapes. These enhancers work together to precisely control the timing and location of gene expression (36, 37). Furthermore, multiple loops facilitate the formation of localized nuclear microenvironments that concentrate transcription factors and coactivators (38). For instance, consistent with previous reports of enhancer-rich regulatory landscapes at the Tbx3-Tbx5-Lhx5 locus in the developing limbs of mouse embryos (39), we found 16 loops for *Tbx3*. Similarly, we found 17 loops for *Tgfb2*, which is in agreement with a report of 15 distinct promoter-interacting regions in human triple-negative breast cancer cells (40). This high degree of regulatory complexity acts as a “regulatory buffer,” whereby enhancer and suppressor loops converge in the regulation of a specific gene. Functionally, this buffering refers to the capacity of redundant regulatory elements to maintain consistent target gene expression levels despite fluctuations or reductions in upstream transcription factors, and environmental variations such as temperature shifts (35, 41, 42). This promotes phenotypic robustness by safeguarding proper developmental outcomes against inevitable environmental stressors or genetic variability (38, 39, 42).

Our finding that “developmental” multi□loop architectures persist in the adult rat frontal cortex supports the idea that these regulatory structures are essential to maintain neuronal identity, connectivity and homeostasis throughout the lifespan. For instance, complex enhancer clusters (e.g., the *Sox2* SRR2-18 cluster) are essential not only for initial development but also for the maintenance of the neural stem and progenitor cell phenotype (43). Given that the mammalian postnatal brain retains diverse neuronal and glial cell types defined by distinct gene expression profiles (44, 45), the high loop density around genes like *Foxo1* and *Spry2* likely implicates the ongoing transcriptional regulation of these genes in the preservation of cell-type specific identity. Furthermore, enhancers regulating proliferation and identity in human neural cells often act on genes associated with neuropsychiatric disorders (e.g., schizophrenia and bipolar disorder) (46), indicating that persistence and the strict control of these regulatory networks within the adult brain are critical for the prevention of cognitive deficits.

To further examine the neural functions related to loop-rich genes, we utilized the text-mining tool GeneCup (47). The results highlighted that several genes are collectively implicated in critical neural processes, including intracellular signaling, transcriptional regulation, neurotransmission and neural maintenance, and are involved in the pathology of neurodegenerative diseases. For example, *Gja1* (Connexin43) is a key driver of astrocyte-specific networks dysregulated in Alzheimer’s disease (48). Astrocyte-specific overexpression of *Gja1* has been found to enhance neuronal viability and mitochondrial recovery in models of traumatic brain injury, highlighting its neuroprotective capacity (49). In the dopaminergic systems, *Tgfb2* variants are associated with susceptibility to Parkinson’s disease, likely due to its role in supporting dopaminergic neuron survival (50). Additionally, *Foxo1* acts as a critical mediator of psychological stress-induced neuroinflammation (51). Elevated *Foxo1* expression was observed in the hippocampus following stress exposure, and its knockdown markedly attenuated stress-induced inflammatory factors in microglia (51, 52). Taken together, the fact that these critical neuroprotective and disease-associated genes are embedded within exceptionally dense chromatin loop networks in our cortical data is highly insightful. It suggests that, much like their functions in other brain regions, the mammalian brain relies on this massive 3D regulatory redundancy to securely buffer the expression level of these genes, thereby ensuring the lifelong functional integrity of the adult cortex against environmental stress and aging.

Although the laboratory rat is an important model for biomedical research, its genomic functional annotation lags behind that of human and mouse (14). This work provides functional data by mapping chromatin interactions across multiple strains in the rat brain. The resulting dataset enhances the utility of the rat as a model organism by helping connect genetic variation to gene regulation. Similar to humans, many disease-associated variants found in rat genetic studies are located in non-coding regions, which complicates the identification of their target genes and functional mechanisms (53, 54). Our data address this challenge by providing physical links between distal regulatory elements and their promoters. This enables a more precise interpretation of non-coding variants by connecting them to specific genes within interacting loop anchors. Furthermore, these data can be integrated with genome-wide association study results using tools like H-MAGMA (55) to prioritize candidate genes. Integrating this chromatin interaction map with transcriptomic and epigenomic data will further enable the prioritization of disease-relevant loci and the inference of their upstream transcriptional regulators (56).

While many of the loops we identified likely mediate P-E interactions, some could connect promoters to silencers, which also have roles in gene regulation (57). Future studies integrating this dataset with other functional genomics data, such as epigenomic maps of active and repressive chromatin marks (e.g., H3K27ac and H3K27me3) and transcriptomic data, will be necessary to further investigate the functional consequences of these 3D interactions.

There are also a few limitations of this study. Consistent with previous Hi-C studies (58), we found that sequencing depth was a primary factor in loop detection, with lower depth yielding fewer, more commonly shared loops between samples. The resolution of detection also influenced the number of loops we identified. Lower resolutions (e.g., 25K) captured a broader set of long-range interactions, whereas higher resolutions (5K) provided more precise boundary definitions but fewer total loops. We thus combined loops from all three resolutions to construct a comprehensive catalog. However, these findings underscore a technical consideration for comparative studies: technical variability can confound the identification of biological differences, such as strain or treatment specific loops. Consequently, our current catalog primarily reflects a consensus regulatory landscape of the rat frontal cortex, rather than strain-to-strain variability. In addition, we generated these data using DNA from the rat frontal cortex, a brain region involved in executive functions and complex behaviors (59). We selected this region for its relevance to the regulatory architecture of many behavioral traits. However, this choice of tissue limits the generalizability of our findings to other tissues, as specific enhancer-promoter loops vary substantially across different cell types (60). It is plausible that other cortical or non-cortical brain areas exhibit alternative chromatin looping patterns.

## METHODS

### Hi-C data generation

Hi-C library construction and sequencing were conducted for the frontal cortex of 10 laboratory rat strains (SHR/OlaIpcv, BN-Lx, BXH6, HXB2, HXB10, HXB23, HXB31, LE/Stm, F344/Stm, and SHR/OlaIpcvxBN/NHsdMcwi) using Arima Hi-C kits, as described previously (21).

### Hi-C reads mapping and loop annotation

We processed the raw sequencing data using the Juicer pipeline (v1.6) (22) and annotated chromatin loops with HiCCUPS (18), implemented at Juicer Tools v1.22.01 in a CUDA 8 environment on Google Colab (https://colab.research.google.com). For two strains, SHR/OlaIpcv (SRR34514260) and HXB10/Ipcv (SRR34514259), SRA submission records under BioProject PRJNA1197090 indicate that each Hi-C sample is represented by a single sample accession but contains two sequencing sets/libraries. SHR/OlaIpcv includes the 592A (S1) and 592B (S2) libraries, whereas HXB10/Ipcv includes the 607B (S3) and 607C (S4) libraries. For HXB10/Ipcv, the two sets were merged and analyzed as a single sample-level Hi-C dataset. In contrast, loop calling and downstream analyses for SHR/OlaIpcv were based on the higher-quality 592B dataset only rather than the merged dataset, because the 592A library showed substantially poorer Hi-C library quality, including a markedly higher PCR duplicate rate (70.8% for 592A versus 11.9% for 592B, where PCR duplicate rate = PCR duplicate read pairs / total sequenced read pairs). HiCCUPS was run at its default resolutions of 5, 10, and 25 kb with Knight–Ruiz (KR) matrix balancing normalization and the --ignore-sparsity flag enabled to ensure consistent loop detection across all chromosomes regardless of local contact density.

### Genome-wide CTCF site prediction

CTCF motifs were curated from multiple publicly available databases (Table S2) including JASPAR 2022 (61), HOCOMOCO v11 (62), SwissRegulon (63), Jolma2013 (64), CTCFBSDB (65), and CIS-BP (ver.3) (66), based on datasets used in the R/Bioconductor data package for CTCF binding in human and mouse (30). Each set of motifs was converted to MEME format using utilities from the MEME Suite (67), including transfac2meme, uniprobe2meme, and jaspar2meme. The resulting MEME files were consolidated into a single MEME-format file. To comprehensively identify all potential CTCF binding sites across the genome, we retained all motifs (Table S2) collected from the databases during FIMO scanning, rather than filtering for redundancy prior to analysis. This approach allows for the capture of subtle variations in motif representations across different databases, thereby enhancing sensitivity. To generate the optional background file for FIMO, we employed the fasta-get-markov utility from the MEME Suite on the mRatBN7.2 genome assembly (command: fasta-get-markov rn7.fa output_background_file.txt). The input MEME file with the background file was scanned against the rat reference genome (mRatBN7.2/rn7) using FIMO (v5.5.4) (23) from the MEME Suite (v5.5.4), installed via Conda (v4.1.0).

### Transcription start sites (TSS), promoter, and gene annotation

TSS annotations were derived from the GTF file (release version of 113) downloaded from ENSEMBL (https://ftp.ensembl.org/pub/release-113/gtf/rattus_norvegicus/Rattus_norvegicus.mRatBN7.2.113.gtf.gz). The initial dataset was subsequently refined by selecting tag and gene_biotype annotated as “Ensembl_canonical” and “protein_coding”, respectively. After removing redundant TSS assigned to the same gene, a final set of 21,725 unique TSS was retained, each corresponding to a distinct gene. Promoter coordinates were obtained from a bed file of Rn_EPDnew_001_rn6.bed (https://epd.expasy.org/ftp/epdnew/R_norvegicus/001/) in the Eukaryotic Promoter Database (EPD) for Rattus norvegicus (68). An initial set of 12,601 promoters based on the Rnor6.0 genome assembly was obtained. These coordinates of the promoters were converted to the mRatBN7.2 (rn7) reference genome using UCSC LiftOver tool (69) with a UCSC Chain file, rn6ToRn7.over.chain.gz (https://hgdownload.soe.ucsc.edu/goldenPath/rn6/liftOver/). Of the 12,533 promoters mapped to rn7, 12,529 unique entries were retained after removing redundant coordinates per gene. We defined the gene’s genomic coordinate based on the coordinates of the last exon (i.e., the exon with the highest rank per transcript). The exon datasets were retrieved from the same GTF file above and from UCSC (https://hgdownload.soe.ucsc.edu/goldenPath/rn7/bigZips/genes/refGene.gtf.gz). This gene coordinate information was subsequently integrated with the TSS and promoter datasets. To associate these promoters with their corresponding gene coordinates, we first mapped EPD promoter identifiers to Ensembl Gene IDs provided by the EPD database using its cross-reference table (https://epd.expasy.org/ftp/epdnew/R_norvegicus/001/db/promoter_ensembl.txt).

### Defining the flanking region of chromatin loops for examining the distribution of CTCF, TSS and promoter across loops

To examine the distribution of genomic features across chromatin loops, we restricted the analysis to loops shorter than 2 megabases (Mb) (18, 24), resulting in the set of 31,019 from an initial non-redundant set of 31,773. We then defined the loop length (L) as the genomic distance between the midpoints of the two anchors in a loop, where each midpoint was calculated as the center between the coordinate pairs (e.g., x1, x2 or y1, y2) of each anchor, with the length of the anchor corresponding to the loop resolution. This definition ensures that L reflects the physical span of the loop between its central anchor positions. Next, we symmetrically extended the region by adding flanking segments of length (L) outward from each of the anchor midpoints. This resulted in a 3L-wide window: the original loop (L) plus two flanking regions (L on each side). These extended windows, which exceeded chromosome boundaries, were excluded (91 loops) because they distort the relative positions of CTCF motifs, TSS, and promoter elements. For the remaining 30,928 loops, we annotated the genomic positions of CTCF motifs, TSS, and promoter regions within the corresponding 3L windows. Their spatial distribution along these normalized windows was then quantified by converting each feature’s coordinate into a relative position using the following transformation:

Relative Position = 3*(Feature coordinate - Loop Start)/L - 1

### Sequential filtering of annotated chromatin loops to define functional interactions

To establish a high-confidence set of functionally relevant chromatin interactions, we applied a multi-step filtering pipeline (Figure 7). We began by aggregating only unique loops identified across the ten samples (27,219 loops removed), followed by filtering out 754 chromatin loops with a genomic distance exceeding 2Mb to ensure the biological relevance of cis-regulatory contacts (70–73), retaining a total of 31,019 loops for downstream analysis.

We then prioritized the loops based on the presence of the architectural protein CTCF, a key regulator of loop formation (28, 74, 75). Using FIMO-predicted CTCF motifs, we quantified the number of unique binding sites within each loop anchor. We retained only loops in which both anchors contained at least 6 CTCF binding sites (Figure 6). Applying this criterion yielded a total of 25,268 loops.

In parallel, we assigned TSS and promoter regions to chromatin loop anchors based on a proximity□based mapping approach. Specifically, we first restricted the search to TSS and promoters located on the anchor□facing side of the loop midpoint, extending outward along the chromosome. After applying this directional constraint, we assigned to each anchor the TSS or promoters with the shortest distance to its midpoint using the distanceToNearest function with the select = “all” option in GenomicRanges R package (76). Then, we applied a multi□stage refinement procedure to obtain a single assignment per anchor for anchors associated with multiple candidate elements due to the “all” option. We removed candidates lacking gene annotations (i.e., candidates with unavailable gene coordinates), prioritized promoter annotations when both promoter and TSS labels were present for the same gene, enforced gene□identity consistency through trimmed gene identifiers, and resolved remaining multi□gene cases using transcript□level features (including exon structure) with additional manual curation to remove low□confidence or uninformative gene symbols (e.g., LOC annotations and Gzmbl2). These steps produced a unique gene assignment for nearly all loop anchors, yielding 61,969 anchors with a single matched gene and three exceptional loops in which both anchors retained two valid assignments (six anchors in total). In sum, this resulted in 61,975 anchor–regulatory element distance measurements. The distance distribution of these finalized assignments was plotted (Figure S8), and 200 kb was selected as the distance threshold.

Next, under the assumption that each chromatin loop corresponds to a single P-E interaction, we removed 13,043 loops in which neither anchor carried a valid TSS or promoter assignment, leaving 17,976 loops with at least one assigned anchor. Among these, 12,707 loops had only one anchor with a valid assignment and were directly retained. For the remaining 5,269 loops in which both anchors carried valid assignments, we selected the anchor whose assigned element was closer to its midpoint. After applying the 200 kb distance threshold to the finalized TSS/promoter-anchor assignments, 17,648 loops were retained.

To obtain a final set of functional loops, we intersected the 17,648 loops from TSS/promoter assignments with the 25,268 loops retained after CTCF motif-based filtering (≥6 binding sites per anchor). This last step yielded 15,085 high-confidence loops jointly supported by the two independent processes (Figure 7).

### Quantification of loop-based interactions in gene-level and functional enrichment analysis

Functional enrichment analysis was performed using g:Profiler (Figure 9) on the top 54 genes (Table S3). These genes were ranked by the highest number of valid loop interactions, ranging from a minimum of 11 to a maximum of 19 interactions per gene.

## ACKNOWLEDGMENTS

We thank Rachel Ward for generating the Hi-C libraries. We gratefully acknowledge the contributions to the Hybrid Rat Diversity Panel (HRDP): Dr. Melinda R. Dwinell (Medical College of Wisconsin) provided the breeders; the Center for Integrative and Translational Genomics at the University of Tennessee Health Science Center (UTHSC) supported colony maintenance; and Mr. Angel Garcia Martinez and Ms. Caroline Jones assisted with breeding. Sequencing data were generated by the UTHSC Molecular Resource Center and University of Tennessee Genomics Core. Computational analyses utilized the University of Tennessee Infrastructure for Scientific Applications and Advanced Computing (ISAAC). This work was supported by grants U01 DA-053672 from NIH/NIDA (to B.M.S., R.W.W., and H.C.).

