## Supplemental file for "Genome-Wide Annotation of Promoter-Enhancer Interactions via Chromatin Loops in the Hybrid Rat Diversity Panel" (from another version of this preprint: version 3, 2026-05-18)

| **Strain** | **Detection Resolution** | | | **Total** |
| --- | --- | --- | --- | --- |
|  | 5K | 10K | 25K |  |
| BN-Lx/Cub | 1491 | 2388 | 2656 | 6535 |
| BXH6/Cub | 1357 | 2406 | 2805 | 6568 |
| F344/Stm | 113 | 808 | 1982 | 2903 |
| HXB10/Ipcv | 1828 | 2574 | 2934 | 7336 |
| HXB2/Ipcv | 521 | 1727 | 2408 | 4656 |
| HXB23/Ipcv | 1976 | 2908 | 2792 | 7676 |
| HXB31/Ipcv | 2849 | 3351 | 2931 | 9131 |
| LE/Stm | 117 | 854 | 2021 | 2992 |
| SHR/OlaIpcv | 856 | 1935 | 2472 | 5263 |
| SHR/OlaIpcv X BN/NHsdMcwi F1 | 969 | 2320 | 2643 | 5932 |

**Table S1. Number of loops by sample and detection resolution**

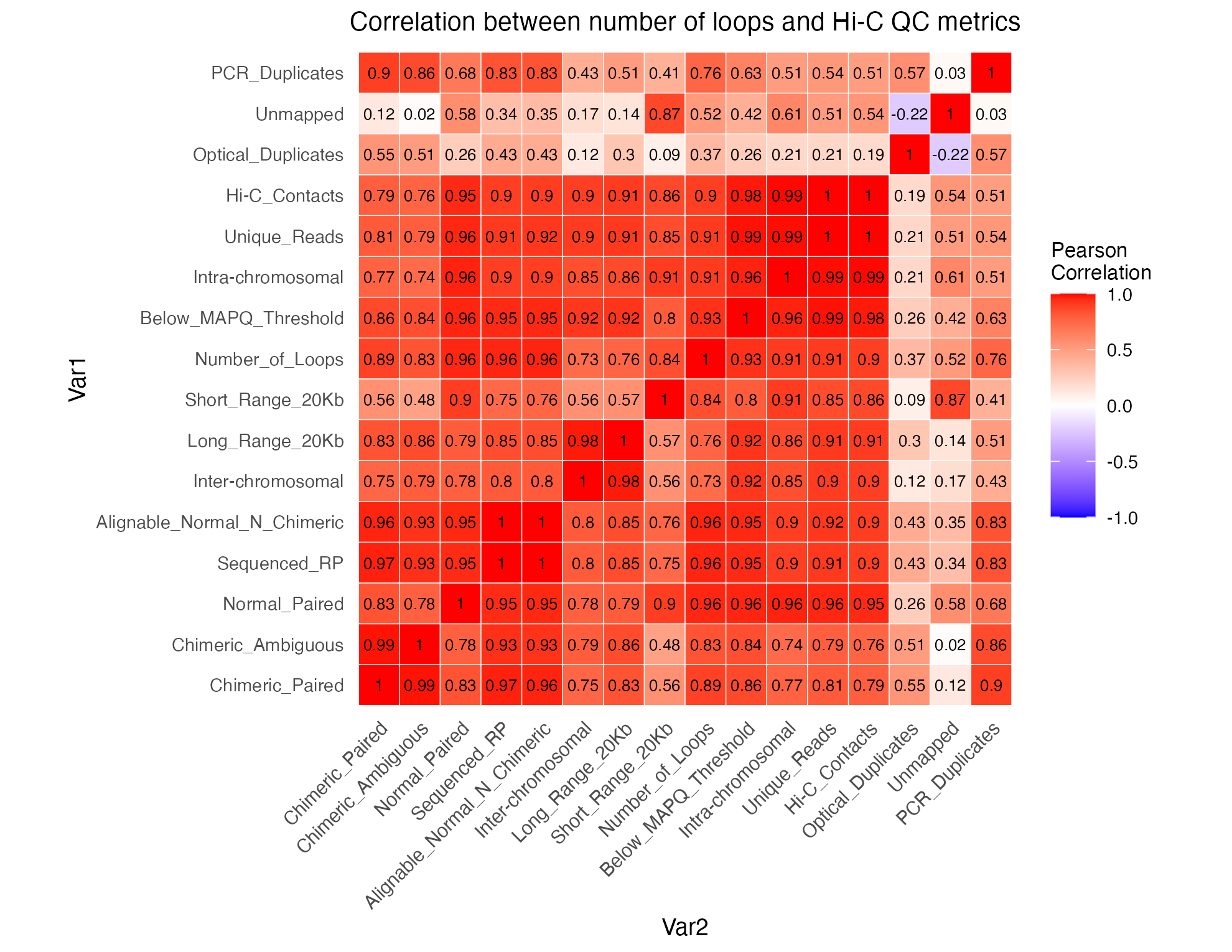
**Figure S1. Pearson correlation between the number of loops and Hi-C sequencing quality control (QC) metrics** Strong positive correlations (in red) were observed between loop count and several key QC features such as the number of Hi-C contacts, unique reads, and alignable non-chimeric reads, suggesting that loop detection is strongly dependent on sequencing depth and data quality. Correlations are symmetric across the diagonal, and the intensity of the color reflects the strength and direction of the relationship.

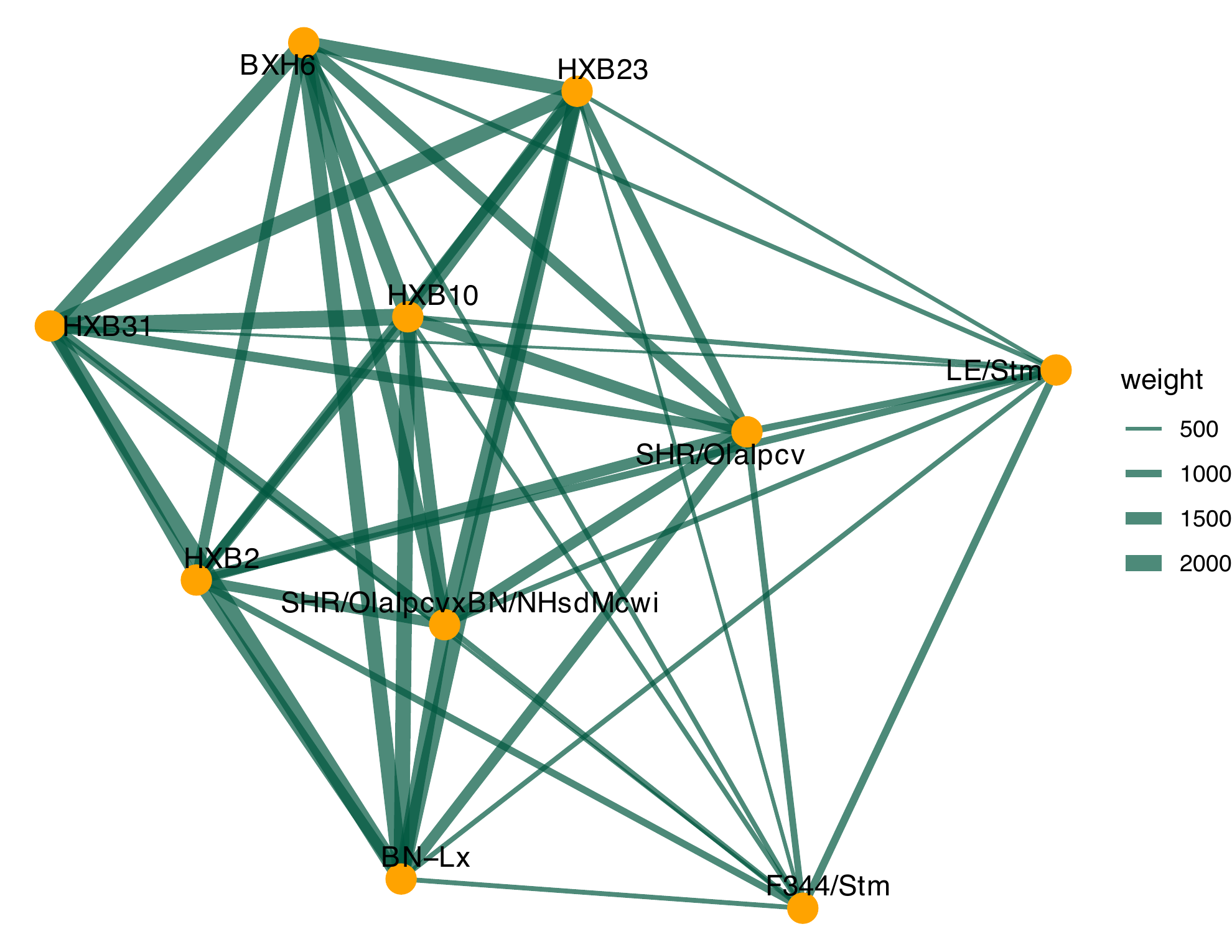

**Figure S2. Shared chromatin loops between rat strains** Each node represents a strain, and the thickness of the connecting edges corresponds to the number of shared loops between strains.

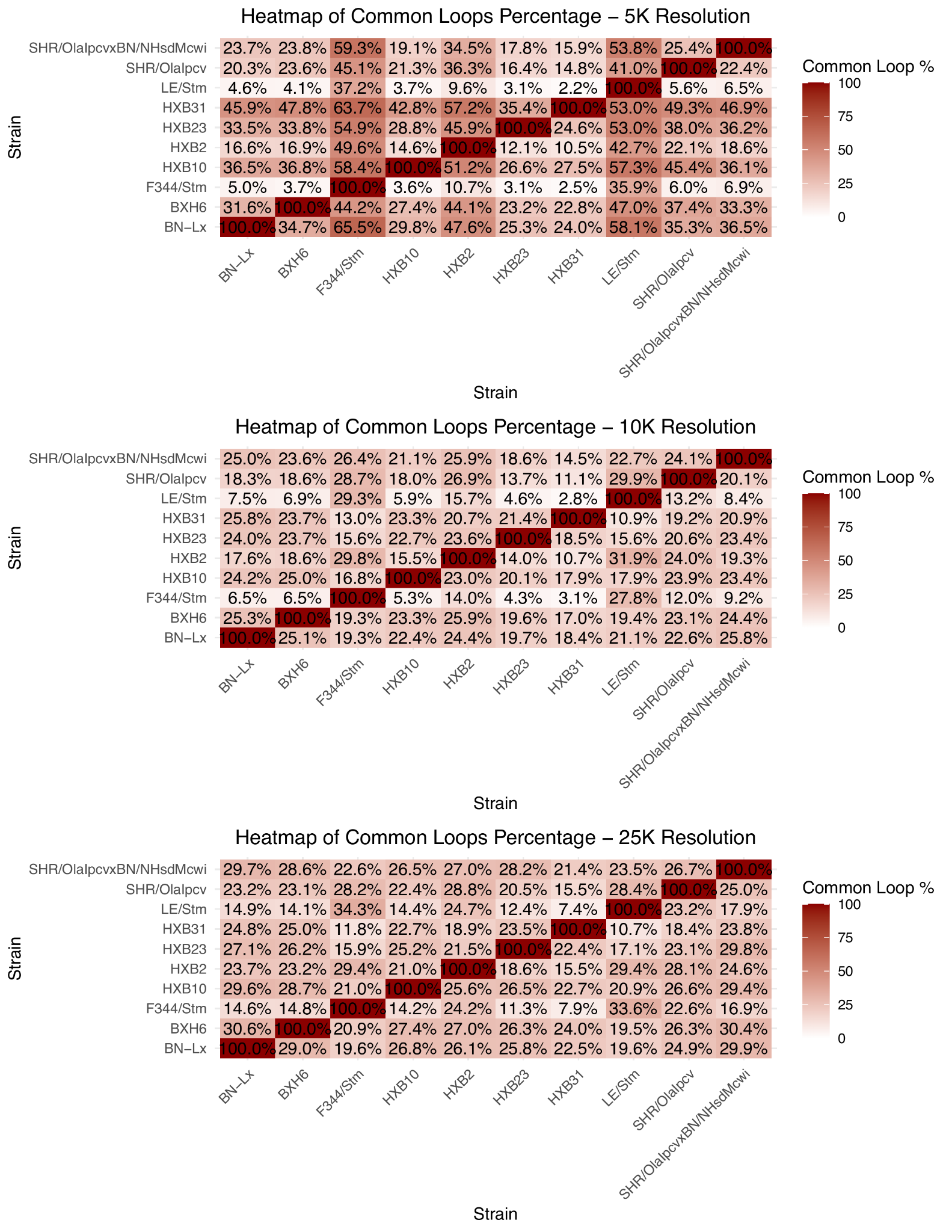

**Figure S3. Heatmaps of pairwise common loop percentages across resolution.** Each cell represents the percentage of loops in strain *i* that are also present in strain *j*. Values are calculated relative to the total number of loops in strain *i*.

| **DB** | **MOTIF** | **WIDTH** | **BEST POSSIBLE MATCH** | **URL** |
| --- | --- | --- | --- | --- |
| **cisBP3.0** | **cisBP_3.0_T094831_3.00\|CTCF\|ENSG00000102974\|M03082_3.00\|C2H2** | **17** | **TACCACTAGGTGGCGCT** | **http://cisbp.ccbr.utoronto.ca/TFreport.php?searchTF=T094831_2.00** |
|  | **cisBP_3.0_T094831_3.00\|CTCF\|ENSG00000102974\|M06045_3.00\|C2H2** | **39** | **CATTACCCCCTCTGGCCACCAGGGGGCGCTGTAGCCCCA** |  |
|  | **cisBP_3.0_T094831_3.00\|CTCF\|ENSG00000102974\|M06233_3.00\|C2H2** | **17** | **TAGGGGGCGCTATTGGG** |  |
|  | **cisBP_3.0_T094831_3.00\|CTCF\|ENSG00000102974\|M07971_3.00\|C2H2** | **15** | **GCCACCAGGGGGCAC** |  |
|  | **cisBP_3.0_T094831_3.00\|CTCF\|ENSG00000102974\|M07972_3.00\|C2H2** | **11** | **AAAAATAAAAA** |  |
|  | **cisBP_3.0_T094831_3.00\|CTCF\|ENSG00000102974\|M07973_3.00\|C2H2** | **19** | **ACACAGGGGATCTCTGTTT** |  |
|  | **cisBP_3.0_T094831_3.00\|CTCF\|ENSG00000102974\|M08320_3.00\|C2H2** | **21** | **GTGGCCACCAGGGGGCGCTGT** |  |
|  | **cisBP_3.0_T094831_3.00\|CTCF\|ENSG00000102974\|M08321_3.00\|C2H2** | **16** | **CCACCAGGGGGCGCCG** |  |
|  | **cisBP_3.0_T094831_3.00\|CTCF\|ENSG00000102974\|M08322_3.00\|C2H2** | **15** | **CCACCAGGGGGCGCC** |  |
|  | **cisBP_3.0_T094831_3.00\|CTCF\|ENSG00000102974\|M08323_3.00\|C2H2** | **16** | **CACCAGGGGGCGCCAG** |  |
|  | **cisBP_3.0_T094831_3.00\|CTCF\|ENSG00000102974\|M08324_3.00\|C2H2** | **15** | **GCCACCAGAGGGCGC** |  |
|  | **cisBP_3.0_T094831_3.00\|CTCF\|ENSG00000102974\|M08325_3.00\|C2H2** | **13** | **GCCACTAGAGGGC** |  |
|  | **cisBP_3.0_T094831_3.00\|CTCF\|ENSG00000102974\|M08326_3.00\|C2H2** | **15** | **GCCACCAGGGGGCGC** |  |
|  | **cisBP_3.0_T094831_3.00\|CTCF\|ENSG00000102974\|M08327_3.00\|C2H2** | **15** | **GCCACCAGGGGGCGC** |  |
|  | **cisBP_3.0_T094831_3.00\|CTCF\|ENSG00000102974\|M08328_3.00\|C2H2** | **15** | **GCCACCAGGGGGCGC** |  |
|  | **cisBP_3.0_T094831_3.00\|CTCF\|ENSG00000102974\|M08329_3.00\|C2H2** | **15** | **CGGCCACCAGGGGGC** |  |
|  | **cisBP_3.0_T094831_3.00\|CTCF\|ENSG00000102974\|M08330_3.00\|C2H2** | **15** | **CCACCAGGGGGCGCC** |  |
|  | **cisBP_3.0_T094831_3.00\|CTCF\|ENSG00000102974\|M08331_3.00\|C2H2** | **21** | **GGCCACCAGGGGGCGCCCGAG** |  |
|  | **cisBP_3.0_T094831_3.00\|CTCF\|ENSG00000102974\|M08332_3.00\|C2H2** | **15** | **CCACCAGGGGGCGCC** |  |
|  | **cisBP_3.0_T094831_3.00\|CTCF\|ENSG00000102974\|M08333_3.00\|C2H2** | **15** | **GCCACCAGGGGGCGC** |  |
|  | **cisBP_3.0_T094831_3.00\|CTCF\|ENSG00000102974\|M08334_3.00\|C2H2** | **15** | **GCCACCAGGGGGCGC** |  |
|  | **cisBP_3.0_T094831_3.00\|CTCF\|ENSG00000102974\|M08335_3.00\|C2H2** | **15** | **CCACCAGGGGGCGCC** |  |
|  | **cisBP_3.0_T094831_3.00\|CTCF\|ENSG00000102974\|M08336_3.00\|C2H2** | **15** | **GCCACCAGGGGGCGC** |  |
|  | **cisBP_3.0_T094831_3.00\|CTCF\|ENSG00000102974\|M08337_3.00\|C2H2** | **15** | **GCCACCAGGGGGCGC** |  |
|  | **cisBP_3.0_T094831_3.00\|CTCF\|ENSG00000102974\|M08338_3.00\|C2H2** | **15** | **TGGCCACCAGGGGGC** |  |
|  | **cisBP_3.0_T094831_3.00\|CTCF\|ENSG00000102974\|M08339_3.00\|C2H2** | **15** | **GGCCACCAGGGGGCG** |  |
|  | **cisBP_3.0_T094831_3.00\|CTCF\|ENSG00000102974\|M08340_3.00\|C2H2** | **12** | **CACCAGGGGGCG** |  |
|  | **cisBP_3.0_T094831_3.00\|CTCF\|ENSG00000102974\|M08341_3.00\|C2H2** | **17** | **GCCACCAGGGGGCGCCA** |  |
|  | **cisBP_3.0_T094831_3.00\|CTCF\|ENSG00000102974\|M08342_3.00\|C2H2** | **15** | **GCCACCAGGGGGCGC** |  |
|  | **cisBP_3.0_T094831_3.00\|CTCF\|ENSG00000102974\|M08343_3.00\|C2H2** | **15** | **GCCACCAGGGGGCGC** |  |
|  | **cisBP_3.0_T094831_3.00\|CTCF\|ENSG00000102974\|M08344_3.00\|C2H2** | **15** | **GCCACCAGGGGGCGC** |  |
|  | **cisBP_3.0_T094831_3.00\|CTCF\|ENSG00000102974\|M08345_3.00\|C2H2** | **18** | **TGGCCACCAGGGGGCGCC** |  |
|  | **cisBP_3.0_T094831_3.00\|CTCF\|ENSG00000102974\|M08346_3.00\|C2H2** | **15** | **GGCCACCAGGGGGCG** |  |
|  | **cisBP_3.0_T094831_3.00\|CTCF\|ENSG00000102974\|M08347_3.00\|C2H2** | **14** | **GCCACCAGGGGGCG** |  |
|  | **cisBP_3.0_T094831_3.00\|CTCF\|ENSG00000102974\|M08348_3.00\|C2H2** | **13** | **CCACCAGAGGGCG** |  |
|  | **cisBP_3.0_T094831_3.00\|CTCF\|ENSG00000102974\|M08349_3.00\|C2H2** | **18** | **TGGCCACCAGGGGGCGCC** |  |
|  | **cisBP_3.0_T094831_3.00\|CTCF\|ENSG00000102974\|M08350_3.00\|C2H2** | **15** | **CCACCAGGGGGCGCT** |  |
|  | **cisBP_3.0_T094831_3.00\|CTCF\|ENSG00000102974\|M08351_3.00\|C2H2** | **15** | **GCCACCAGGGGGCAC** |  |
|  | **cisBP_3.0_T094831_3.00\|CTCF\|ENSG00000102974\|M08352_3.00\|C2H2** | **17** | **CCACCAGGGGGCGCCGG** |  |
|  | **cisBP_3.0_T094831_3.00\|CTCF\|ENSG00000102974\|M08353_3.00\|C2H2** | **18** | **GCCACCAGGGGGCGCTAT** |  |
|  | **cisBP_3.0_T094831_3.00\|CTCF\|ENSG00000102974\|M08354_3.00\|C2H2** | **15** | **CCACCAGGGGGCGCT** |  |
|  | **cisBP_3.0_T094831_3.00\|CTCF\|ENSG00000102974\|M08355_3.00\|C2H2** | **15** | **CCACTAGGTGGCACT** |  |
|  | **cisBP_3.0_T094831_3.00\|CTCF\|ENSG00000102974\|M08356_3.00\|C2H2** | **14** | **GCCACCAGGGGGCG** |  |
|  | **cisBP_3.0_T094831_3.00\|CTCF\|ENSG00000102974\|M08357_3.00\|C2H2** | **20** | **TGGCCACCAGGGGGCGCTAT** |  |
|  | **cisBP_3.0_T094831_3.00\|CTCF\|ENSG00000102974\|M08358_3.00\|C2H2** | **17** | **CCACCAGGGGGCGCCAG** |  |
|  | **cisBP_3.0_T094831_3.00\|CTCF\|ENSG00000102974\|M08359_3.00\|C2H2** | **11** | **CCACCAGAGGG** |  |
|  | **cisBP_3.0_T094831_3.00\|CTCF\|ENSG00000102974\|M08360_3.00\|C2H2** | **15** | **CCACCAGGGGGCGCT** |  |
|  | **cisBP_3.0_T094831_3.00\|CTCF\|ENSG00000102974\|M08361_3.00\|C2H2** | **13** | **CCACCAGGGGGCG** |  |
|  | **cisBP_3.0_T094831_3.00\|CTCF\|ENSG00000102974\|M08362_3.00\|C2H2** | **15** | **GCCACCAGGGGGCGC** |  |
|  | **cisBP_3.0_T094831_3.00\|CTCF\|ENSG00000102974\|M08363_3.00\|C2H2** | **12** | **CACCAGGGGGCG** |  |
|  | **cisBP_3.0_T094831_3.00\|CTCF\|ENSG00000102974\|M08364_3.00\|C2H2** | **18** | **CCACCAGAGGGCGCTATA** |  |
|  | **cisBP_3.0_T094831_3.00\|CTCF\|ENSG00000102974\|M08365_3.00\|C2H2** | **18** | **GGCCACCAGGGGGCGCTA** |  |
|  | **cisBP_3.0_T094831_3.00\|CTCF\|ENSG00000102974\|M08366_3.00\|C2H2** | **21** | **CGGCCACCAGGGGGCGCCGGG** |  |
|  | **cisBP_3.0_T094831_3.00\|CTCF\|ENSG00000102974\|M08367_3.00\|C2H2** | **15** | **AGTGCCATCTAGTGG** |  |
|  | **cisBP_3.0_T094831_3.00\|CTCF\|ENSG00000102974\|M08368_3.00\|C2H2** | **20** | **GCCACCAGGGGGCGCCGGAG** |  |
|  | **cisBP_3.0_T094831_3.00\|CTCF\|ENSG00000102974\|M08369_3.00\|C2H2** | **18** | **TGGCCACCAGGGGGCGCC** |  |
|  | **cisBP_3.0_T094831_3.00\|CTCF\|ENSG00000102974\|M08370_3.00\|C2H2** | **15** | **GCCACCAGGGGGCGC** |  |
|  | **cisBP_3.0_T094831_3.00\|CTCF\|ENSG00000102974\|M08371_3.00\|C2H2** | **15** | **GCCACCAGGGGGCGC** |  |
|  | **cisBP_3.0_T094831_3.00\|CTCF\|ENSG00000102974\|M08372_3.00\|C2H2** | **15** | **GCCACCAGGGGGCGC** |  |
|  | **cisBP_3.0_T094831_3.00\|CTCF\|ENSG00000102974\|M08373_3.00\|C2H2** | **21** | **TTTGGCCACTAGGGGGCACTA** |  |
|  | **cisBP_3.0_T094831_3.00\|CTCF\|ENSG00000102974\|M08374_3.00\|C2H2** | **17** | **GCCACCAGATGGCACTA** |  |
|  | **cisBP_3.0_T094831_3.00\|CTCF\|ENSG00000102974\|M08375_3.00\|C2H2** | **15** | **GCCACCAGGGGGCGC** |  |
|  | **cisBP_3.0_T094831_3.00\|CTCF\|ENSG00000102974\|M08376_3.00\|C2H2** | **14** | **CCACCAGGGGGCGC** |  |
|  | **cisBP_3.0_T094831_3.00\|CTCF\|ENSG00000102974\|M08377_3.00\|C2H2** | **15** | **CCACCAGGGGGCGCT** |  |
|  | **cisBP_3.0_T094831_3.00\|CTCF\|ENSG00000102974\|M08378_3.00\|C2H2** | **18** | **TGGCCACCAGGGGGCGCT** |  |
|  | **cisBP_3.0_T094831_3.00\|CTCF\|ENSG00000102974\|M08379_3.00\|C2H2** | **15** | **GGCCACCAGGGGGCG** |  |
|  | **cisBP_3.0_T094831_3.00\|CTCF\|ENSG00000102974\|M08380_3.00\|C2H2** | **13** | **CCACCAGGGGGCG** |  |
|  | **cisBP_3.0_T094831_3.00\|CTCF\|ENSG00000102974\|M08381_3.00\|C2H2** | **15** | **GGCCACCAGGGGGCG** |  |
|  | **cisBP_3.0_T094831_3.00\|CTCF\|ENSG00000102974\|M08382_3.00\|C2H2** | **15** | **CCACCAGGGGGCGCC** |  |
|  | **cisBP_3.0_T094831_3.00\|CTCF\|ENSG00000102974\|M08383_3.00\|C2H2** | **15** | **CCACCAGGGGGCGCC** |  |
|  | **cisBP_3.0_T094831_3.00\|CTCF\|ENSG00000102974\|M08734_3.00\|C2H2** | **15** | **GCCACCAGGGGGCGC** |  |
|  | **cisBP_3.0_T094831_3.00\|CTCF\|ENSG00000102974\|M08735_3.00\|C2H2** | **31** | **CTGCAGTTCCAGCTGTGGCCACCAGGGGGCG** |  |
|  | **cisBP_3.0_T094831_3.00\|CTCF\|ENSG00000102974\|M08736_3.00\|C2H2** | **33** | **CTGCAGTTCCCCCTTCTGGCCACTAGGGGGCGC** |  |
|  | **cisBP_3.0_T094831_3.00\|CTCF\|ENSG00000102974\|M09031_3.00\|C2H2** | **15** | **GCCACCAGGGGGCGC** |  |
|  | **cisBP_3.0_T094831_3.00\|CTCF\|ENSG00000102974\|M09072_3.00\|C2H2** | **23** | **GGCCACCAGGGGGCAGCATAGCC** |  |
|  | **cisBP_3.0_T094831_3.00\|CTCF\|ENSG00000102974\|M09073_3.00\|C2H2** | **23** | **GGCCACCAGGGGGCGCCAGAGAC** |  |
|  | **cisBP_3.0_T094831_3.00\|CTCF\|ENSG00000102974\|M09074_3.00\|C2H2** | **22** | **GGCCACCAGGGGGCGCTGTGGC** |  |
|  | **cisBP_3.0_T094831_3.00\|CTCF\|ENSG00000102974\|M09075_3.00\|C2H2** | **21** | **GGCCACTAGGTGGCGCTGTTG** |  |
|  | **cisBP_3.0_T094831_3.00\|CTCF\|ENSG00000102974\|M09076_3.00\|C2H2** | **16** | **GGCCACCAGGGGGCGC** |  |
|  | **cisBP_3.0_T094831_3.00\|CTCF\|ENSG00000102974\|M09077_3.00\|C2H2** | **20** | **CACCAGCAGGGGGCGCGCGG** |  |
|  | **cisBP_3.0_T094831_3.00\|CTCF\|ENSG00000102974\|M09078_3.00\|C2H2** | **26** | **TTCCACATCTGGCCACCAGGGGGCGC** |  |
|  | **cisBP_3.0_T094831_3.00\|CTCF\|ENSG00000102974\|M09424_3.00\|C2H2** | **15** | **CCAGCAGGGGGCGCT** |  |
|  | **cisBP_3.0_T094831_3.00\|CTCF\|ENSG00000102974\|M09992_3.00\|C2H2** | **19** | **TGGCCACCAGGGGGCGCCA** |  |
|  | **cisBP_3.0_T094831_3.00\|CTCF\|ENSG00000102974\|M10626_3.00\|C2H2** | **20** | **TGGCCACCAGGTGGCACTAT** |  |
|  | **cisBP_3.0_T094831_3.00\|CTCF\|ENSG00000102974\|M10627_3.00\|C2H2** | **20** | **TGGCCACTGTTGGAACTGCA** |  |
|  | **cisBP_3.0_T100985_3.00\|Ctcf\|ENSMUSG00000005698\|M08605_3.00\|C2H2** | **15** | **CCACCAGGGGGCGCT** |  |
|  | **cisBP_3.0_T100985_3.00\|Ctcf\|ENSMUSG00000005698\|M10089_3.00\|C2H2** | **20** | **TGGCCACCAGGGGGCGCTGG** |  |
| **ctcfbsdb** | **ctcf_uthsc_EMBL_M1** | **14** | **CGCCCCCTGGTGGC** | **https://insulatordb.uthsc.edu/download/CTCFBSDB_PWM.mat** |
|  | **ctcf_uthsc_EMBL_M2** | **9** | **GGAACTGCA** |  |
|  | **ctcf_uthsc_REN_20** | **20** | **GTGGCCACCAGGGGGCGCCC** |  |
|  | **ctcf_uthsc_MIT_LM2** | **19** | **TGACCACTAGATGGCAGTA** |  |
|  | **ctcf_uthsc_MIT_LM7** | **20** | **TGGCCAGCAGATGGCGCTGT** |  |
|  | **ctcf_uthsc_MIT_LM23** | **20** | **TGACCACCAGGTGGTGCTGT** |  |
| **hocomoco11** | **CTCF_HUMAN.H11MO.0.A_YNRCCRSYAGRKGGCRSYV** | **19** | **TGGCCACCAGGGGGCGCCA** | **http://hocomoco.autosome.ru/motif/CTCF_HUMAN.H11MO.0.A** |
|  | **CTCF_MOUSE.H11MO.0.A_YNRCCRSYAGRKGGCRSYVB** | **20** | **TGGCCACCAGGGGGCGCTGG** | **http://hocomoco.autosome.ru/motif/CTCF_MOUSE.H11MO.0.A** |
| **jaspar2022** | **MA0139.1_MA0139.1** | **19** | **TGGCCACCAGGGGGCGCTA** | **http://jaspar.genereg.net/matrix/MA0139.1** |
|  | **MA1929.1_MA1929.1** | **34** | **CTGCAGTTCCAGCTGTGGCCACCAGGGGGCGCTA** | **http://jaspar.genereg.net/matrix/MA1929.1** |
|  | **MA1930.1_MA1930.1** | **35** | **CTGCAGTTCCCCCTTCTGGCCACTAGGGGGCGCTA** | **http://jaspar.genereg.net/matrix/MA1930.1** |
| **jolma2013** | **5194.motif_CTCF** | **17** | **AGCGCCACCTAGTGGTA** | **https://footprintdb.eead.csic.es//index.php?motif=347a14880822fa09a386e91ac9acf6e7** |
| **swissregulon** | **ctcf.p2_hg18_swissregulon_ctcf.p2_hg18_for_jaspar2meme** | **20** | **TTGGCCACCAGGGGGCACTA** | **https://swissregulon.unibas.ch/wm/?wm=CTCF.p2&org=hg18** |

**Table S2. CTCF motif models used for genome-wide FIMO scanning.**

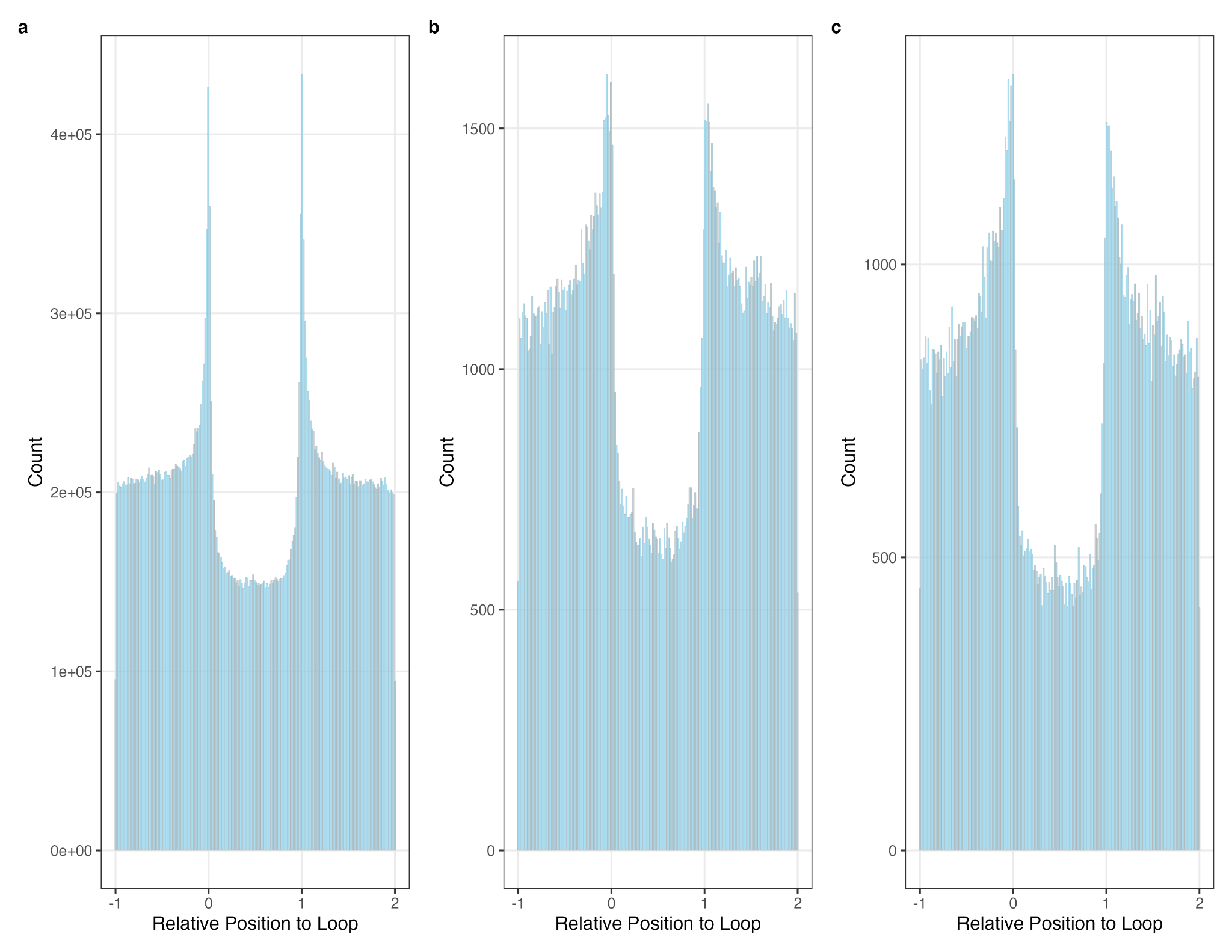

**Figure S4. Histograms of positional distribution for three genomic components: CTCF transcription start sites (TSS), and promoters.** Bar plots depict the count-based distribution of each component relative to chromatin loops, spanning ±1 loop length. Each panel (a–c) visualizes how CTCF, TSS, and promoter elements are spatially arranged around loop anchors. All histograms were constructed using 200 bins. **a.** CTCF binding sites show two prominent and symmetric peaks at both loop anchors, indicating strong and consistent enrichment at loop boundaries. **b.** TSS distribution reveals enrichment is present near loop anchors. Multiple minor peaks suggest increased positional variability or noise. **c.** Promoter counts also exhibit increased signal near each anchor at similar relative distances even though the signals are less defined and accompanied by more dispersed noise peaks compared to CTCF.

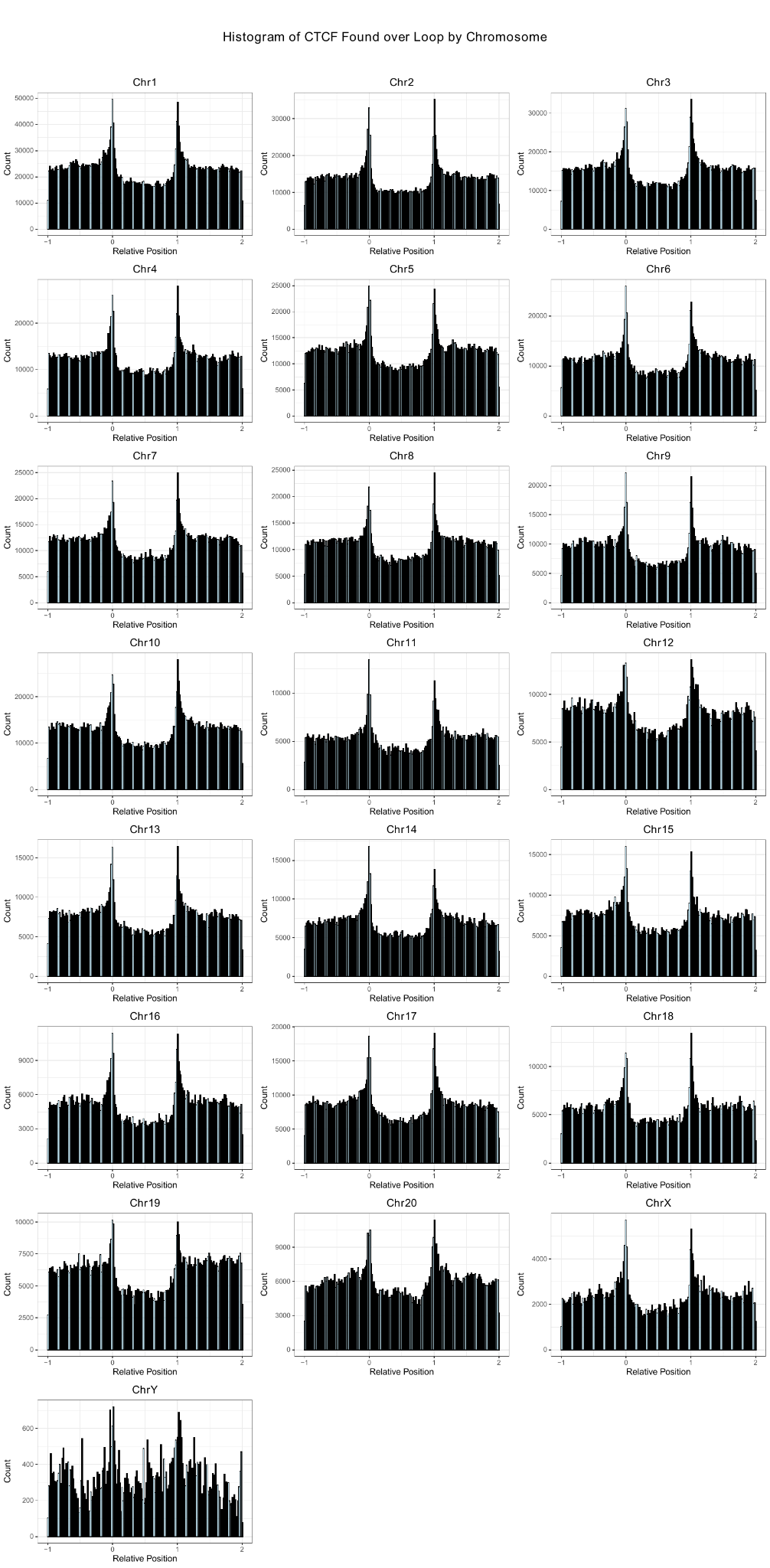
**Figure S5. Distribution of CTCF across chromosomes.** Loop lengths were normalized to 1 unit. The histogram contains all loops detected from 10 samples.

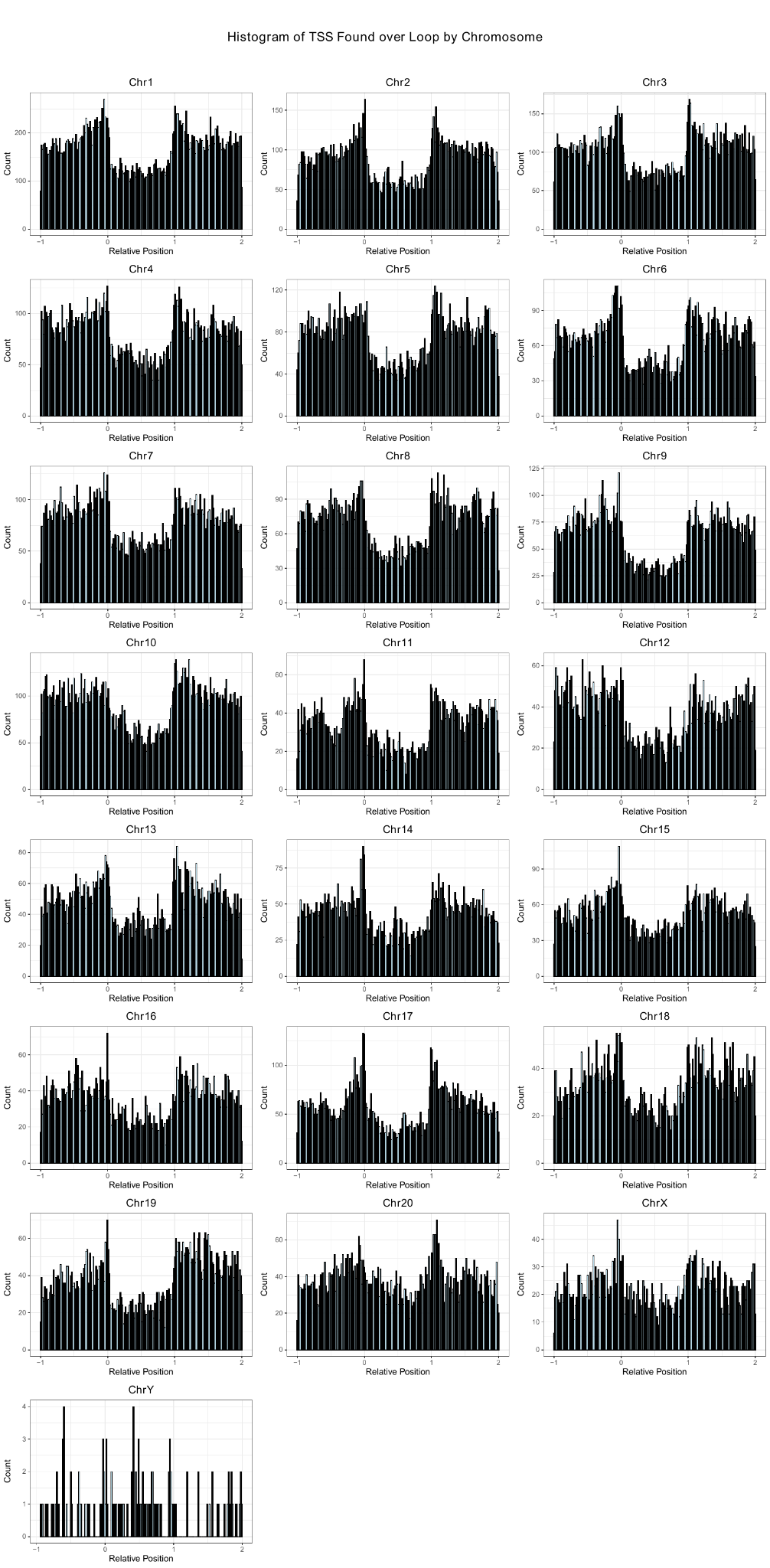
**Figure S6. Distribution of TSS across chromosomes.** Loop lengths were normalized to 1 unit. The histogram contains loops detected from all 10 samples.

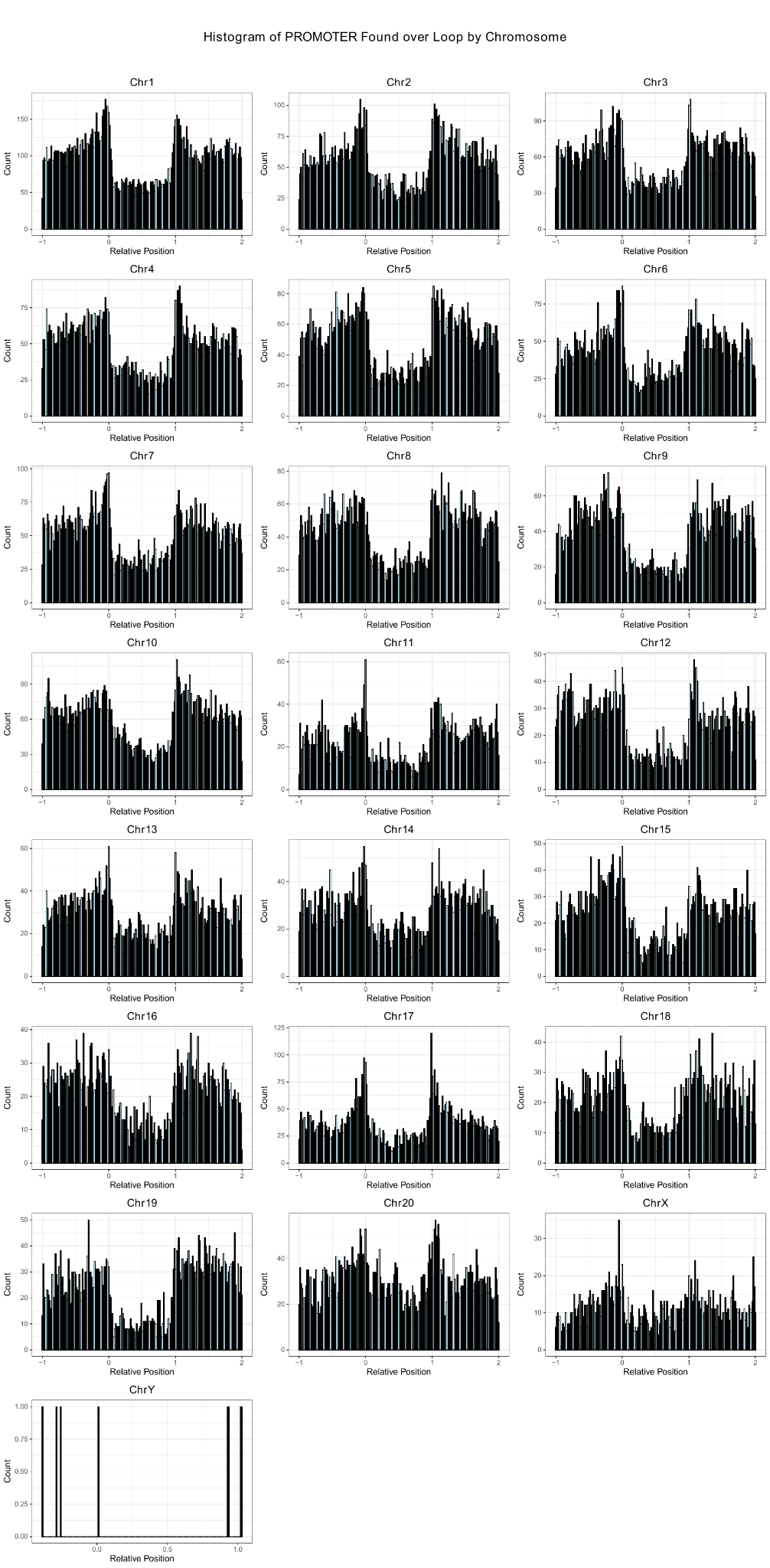
**Figure S7. Distribution of promoters across chromosomes.** Loop lengths were normalized to 1 unit. The histogram contains loops detected from all 10 samples.

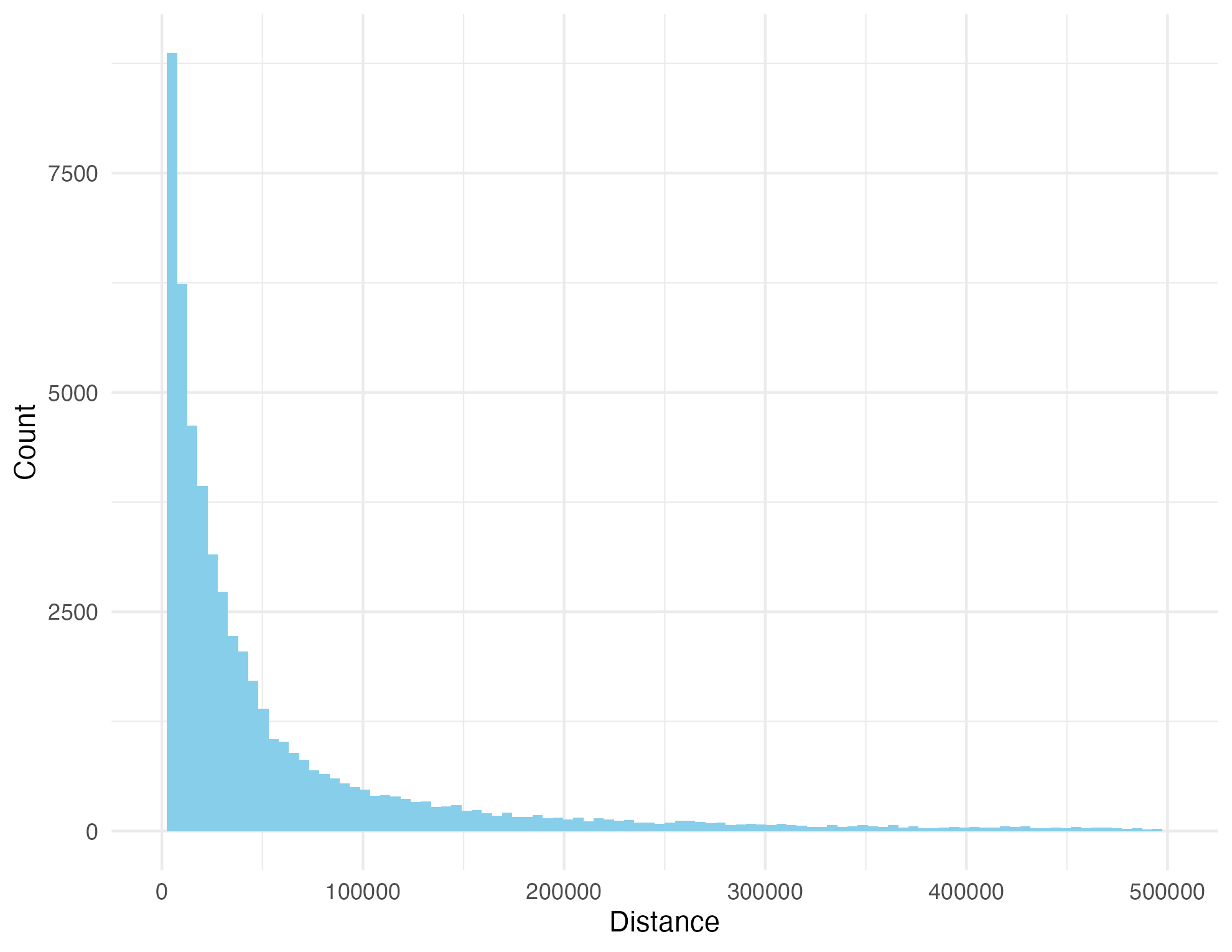

**Figure S8**. Distribution of distances between TSS/Promoter and target anchor

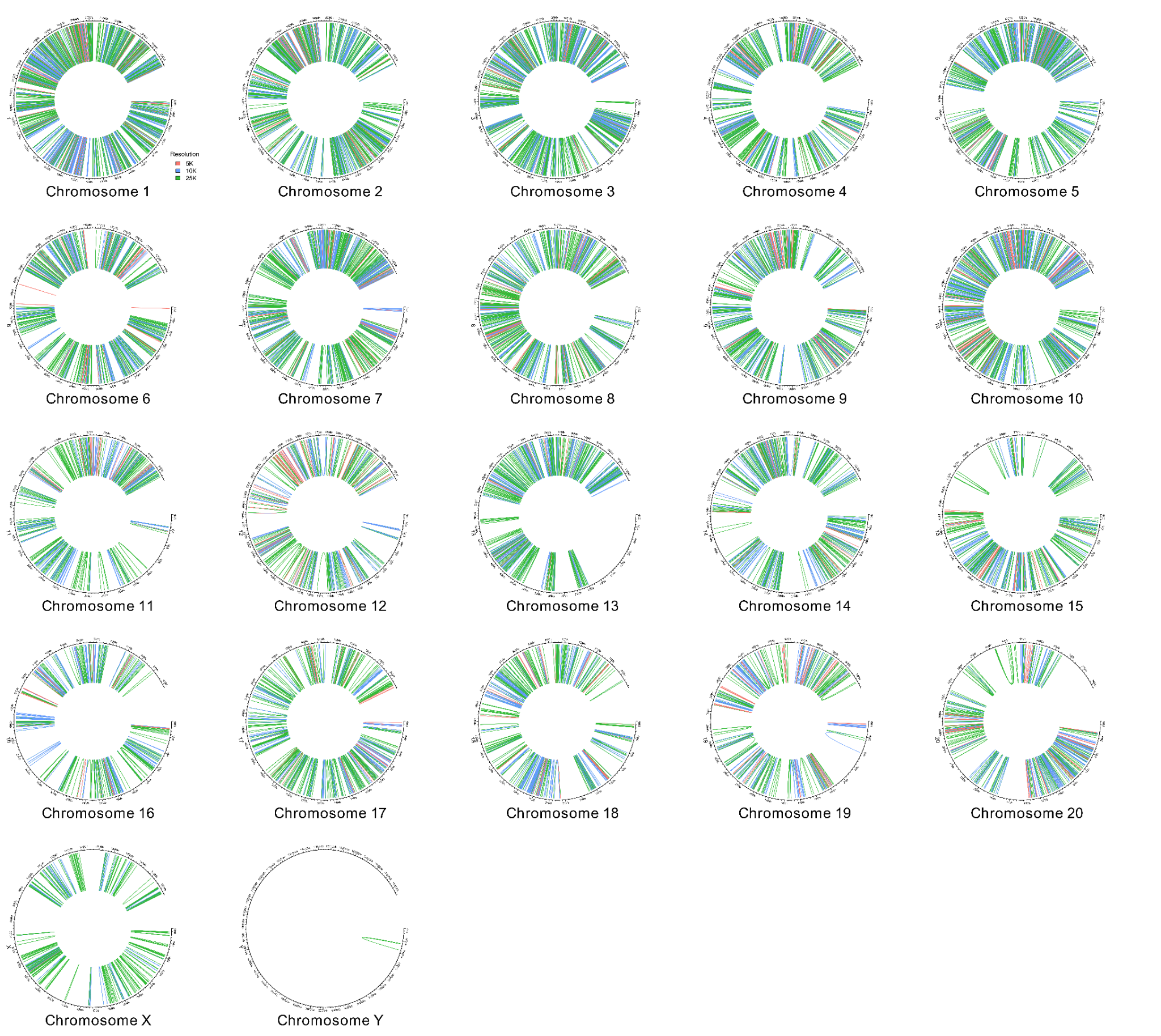

**Figure S9. Circos plot for the functional loops.**

| No. | Gene | Number of interactions |
| --- | --- | --- |
| 1 | Foxo1 | 19 |
| 2 | Gja1 | 19 |
| 3 | Spry2 | 19 |
| 4 | Tgfb2 | 17 |
| 5 | Nfil3 | 16 |
| 6 | Sgk1 | 16 |
| 7 | Tbx3 | 16 |
| 8 | Tsc22d1 | 16 |
| 9 | Foxc1 | 15 |
| 10 | Ptpru | 15 |
| 11 | Arl4c | 14 |
| 12 | Gnas | 14 |
| 13 | LOC100910078 | 14 |
| 14 | LOC108353098 | 14 |
| 15 | Plk2 | 14 |
| 16 | AABR07043557.1 | 13 |
| 17 | En1 | 13 |
| 18 | Foxf2 | 13 |
| 19 | Plpp3 | 13 |
| 20 | Tmem161b | 13 |
| 21 | Tmem200c | 13 |
| 22 | Trmt9b | 13 |
| 23 | Wnt7b | 13 |
| 24 | Ypel2 | 13 |
| 25 | Zfp36l2 | 13 |
| 26 | Cep128 | 12 |
| 27 | Col12a1 | 12 |
| 28 | Dgkg | 12 |
| 29 | Fat4 | 12 |
| 30 | Irf2bp2 | 12 |
| 31 | Kctd12 | 12 |
| 32 | LOC102556715 | 12 |
| 33 | Net1 | 12 |
| 34 | Pdzrn4 | 12 |
| 35 | Sdc3 | 12 |
| 36 | Tmem236 | 12 |
| 37 | Aplnr | 11 |
| 38 | Basp1 | 11 |
| 39 | Cd247 | 11 |
| 40 | Col22a1 | 11 |
| 41 | Dnaaf11 | 11 |
| 42 | Ebf1 | 11 |
| 43 | Efnb1 | 11 |
| 44 | Enc1 | 11 |
| 45 | Fjx1 | 11 |
| 46 | Gprc5b | 11 |
| 47 | Ifit1bl | 11 |
| 48 | Nfkbia | 11 |
| 49 | Pgm5 | 11 |
| 50 | Rpl14l1 | 11 |
| 51 | Rtca | 11 |
| 52 | Six3 | 11 |
| 53 | Tnfsf8 | 11 |
| 54 | Zfp36l1 | 11 |

**Table S3** This table lists genes that exhibit multiple valid interactions. Genes with the same number of interactions are ordered alphabetically.
